# Photochemical probe identification of the small-molecule binding site in a mammalian membrane-bound *O*-acyltransferase

**DOI:** 10.1101/2020.10.16.342477

**Authors:** Thomas Lanyon-Hogg, Markus Ritzefeld, Leran Zhang, Balazs Pogranyi, Milon Mondal, Lea Sefer, Callum D. Johnston, Claire E. Coupland, Sebastian A. Andrei, Joshua Newington, Anthony I. Magee, Christian Siebold, Edward W. Tate

**Affiliations:** Department of Pharmacology, University of Oxford, Oxford, OX1 3QT, UK; Department of Chemistry, Imperial College London, London, W12 0BZ, UK; Division of Structural Biology, Wellcome Centre for Human Genetics, University of Oxford, Oxford, OX3 7BN, UK; CardioRespiratory Interface Section, National Heart & Lung Institute, Imperial College London, London, SW7 2AZ, UK

**Keywords:** Enzymes, Hedgehog acyltransferase, Hedgehog signalling, Medicinal chemistry, Membrane-bound *O*-acyltransferase, Photoaffinity labelling

## Abstract

The mammalian membrane-bound *O*-acyltransferase (MBOAT) superfamily is involved in biological processes including growth, development and appetite sensing. MBOATs are attractive drug targets in cancer and obesity; however, information on the binding site and molecular mechanisms underlying small-molecule inhibition is elusive. This study reports development of a photochemical probe to interrogate the small-molecule binding site in the human MBOAT Hedgehog acyltransferase (HHAT) based on HHAT inhibitor RUSKI-201. Structure-activity relationship investigation identified the improved enantiomeric inhibitor **IMP-1575**, which is the most potent HHAT inhibitor reported to-date, and guided rational design of a photocrosslinking probe that maintained HHAT-inhibitory potency. Photocrosslinking and proteomic sequencing of HHAT delivered identification of the first small-molecule binding site in a mammalian MBOAT. Topology and homology data suggested a potential mechanism for HHAT inhibition which was confirmed via kinetic analysis. Our results provide an optimal HHAT inhibitor **IMP-1575** (*K*_i_ = 38 nM) and a strategy for mapping of interaction sites in MBOATs.

---

Members of the membrane-bound *O*-acyltransferase (MBOAT) superfamily of proteins are involved in several critically important biological pathways.^[1]^ In humans, these include Wnt acyltransferase Porcupine (PORCN),^[2]^ Hedgehog acyltransferase (HHAT)^[3]^ and ghrelin *O*-acyltransferase (GOAT)^[4]^ which regulate Wnt and Hedgehog signaling, and appetite sensing, respectively. These MBOATs are therefore attractive therapeutic targets in cancer and obesity.^[1]^ Structural information for mammalian MBOATs is highly sought after, and recent years have seen several key advances. The membrane topology of various mammalian MBOATs has been experimentally determined, supporting a conserved arrangement of multiple transmembrane helices and catalytic residues.^[1a]^ Recent determination of the structure of a bacterial MBOAT, DltB, provided the first insights into MBOAT architecture and mechanism,^[5]^ revealing that DltB forms a pore-like structure with the active site in the centre of the pore, and providing a rationale for how MBOATs can combine substrates present on opposite sides of biological membranes.^[5]^ *De novo* computational predictions suggest GOAT may adopt a similar structure,^[6]^ and the first cryo-EM structures of a human MBOAT, diacylglycerol *O*-acyltransferase 1 (DGAT1),^[7]^ confirmed the pore-like architecture of MBOATs and provided insights into the catalytic mechanism. However, the nature of small-molecule MBOAT inhibition remains largely unknown.

Hedgehog signaling drives growth during development and is reactivated in certain cancers, making HHAT an attractive therapeutic target to block aberrant signaling.^[1]^ HHAT *N*-acylates Hedgehog signaling proteins with palmitic (C16:0) fatty acid, using palmitoyl-Coenzyme A (Pal-CoA) as the lipid donor substrate (Figure 1A).^[3]^ *N*-acylation of Sonic Hedgehog protein (SHH) by HHAT and C-terminal auto-*O*-cholesterylation^[8]^ is required for signaling function. We previously determined the membrane topology of HHAT, which consists of ten transmembrane helices, two reentrant loops, and one palmitoylation-tethered reentrant loop (Figure 1B).^[9]^ Remarkably these experimental analyses undertaken in live cells placed the MBOAT signature residues His379 and Asp339 on opposite sides of the membrane, raising questions regarding the potential role of these residues in the catalytic mechanism. There is one known series of small-molecule inhibitors of HHAT (termed ‘RUSKI’), with RUSKI-201 (**1**, Figure 1C) the only molecule in this class with proven on-target cellular activity over a non-cytotoxic concentration range.^[10]^ **1** contains two undefined stereocentres, with further scope for optimization of the structure-activity relationship (SAR) to generate higher potency inhibitors. There is also currently no information about this series’ binding mode at HHAT, or indeed the binding site of any inhibitor of a mammalian MBOAT. Photocrosslinking of ligands to proteins followed by proteomic mass spectrometry-based sequencing is a powerful method to identify binding sites.^[11]^ However, such studies remain technically challenging for integral membrane proteins.^[12]^ Here, we report investigation of the RUSKI-201 SAR leading to rational design of a photochemical probe that successfully identified the small-molecule binding site in HHAT for the first time, providing insights into the mechanism of HHAT inhibition.

**Figure 1.**
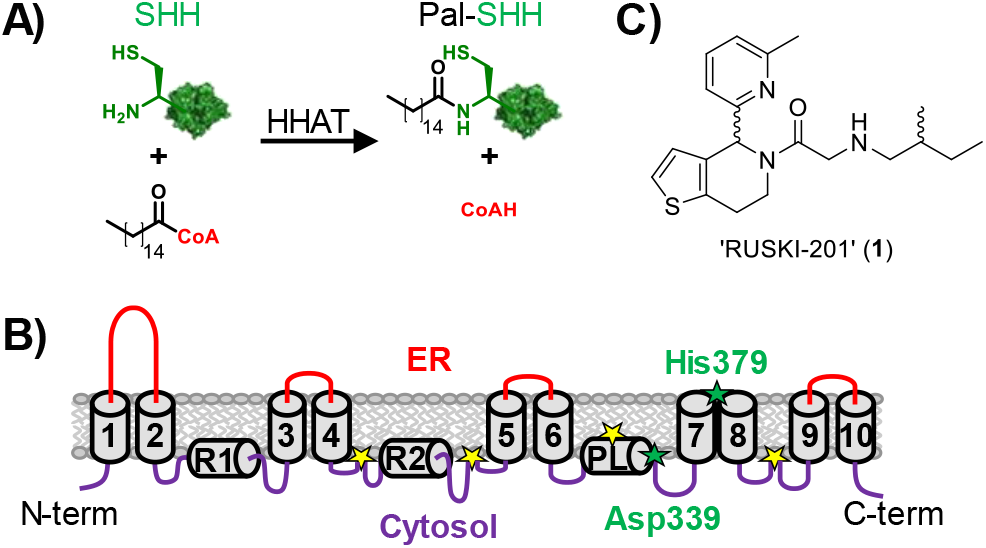
Hedgehog acyltransferase (HHAT) function and topology. A) *N*-acylation reaction of Sonic Hedgehog (SHH) with palmitoyl-CoA catalysed by HHAT. B) Experimentally determined topology model for HHAT,^[9]^ showing ten transmembrane loops (1-10), two re-entrant loops (R1-2) and one palmitoylation-tethered loop (PL). Yellow stars: sites of palmitoylation; green stars: signature MBOAT residues involved in catalysis. C) Structure of RUSKI-201, the only previously known highly selective HHAT inhibitor.^[10]^

The amide-linked side chain of **1** was selected for SAR investigation. The 4,5,6,7-tetrahydrothieno[3,2-*c*]pyridine core (**2**) was synthesized via Bischler–Napieralski cyclization,^[13]^ or via a shorter and higher yielding Pictet-Spengler cyclization (Scheme S1). Acetylated derivative **3** was prepared to investigate the importance of the aminoalkyl chain. Amide-linked sidechain derivatives **4**-**6** were prepared via *N*-benzyl-protected intermediates to enhance handling of the amines by decreasing volatility (Scheme S2). **1** contains two undefined stereocentres, in the central ring system and in the 2-methylbutylamino chain. To investigate the stereochemical requirements for inhibition, (*S*)-2-methylbutylamine was used to prepare **4** with defined sidechain stereochemistry. 3-methylbutylamine (**5**) and 2-methylpropylamine (**6**) derivatives were prepared to remove the stereocentre. To investigate the importance of the sidechain secondary amine, tertiary amine **7** and cyclic derivative **8** were prepared from *N*-methyl isobutylamine and 3-methyl piperidine, respectively. To further probe the secondary amine, urea derivative **9** was prepared from isopentylamine (Scheme S2).

We recently reported the acylation-coupled lipophilic induction of polarization (Acyl-cLIP) assay, a facile and universally applicable method to monitor any enzyme processing lipid post-translational modifications which uses the hydrophobicity increase on lipidation to drive a polarized fluorescence readout.^[14]^ Dose-response analysis of HHAT inhibition was conducted using real-time Acyl-cLIP analysis (Table 1). **1** exhibited a half-maximal inhibitory concentration (IC_50_) of 2.0 μM (95% confidence interval (CI) = 1.4-2.8 μM).^[14]^ Core **2** showed no inhibition, whereas acetamide **3** showed reduced activity (IC_50_ = 13 μM, 95% CI = 7.4-24 μM), indicating the amide is required for activity and that the aminoalkyl chain improves potency. *S*-enantiomer **4** exhibited ^~^2-fold reduced potency (IC_50_ = 3.7 μM, 95% CI = 1.7-8.2 μM) compared to **1**, indicating stereochemistry at this position minimally impacts activity. Moving the methyl group by one position in **5** showed a minor decrease in potency (IC_50_ = 4.6 μM, 95% CI = 3.1-6.9 μM), whereas shortening the alkyl chain **6** increased potency modestly compared to **1** (IC_50_ = 1.3 μM, 95% CI = 0.88-2.0 μM) whilst also removing the stereocentre. Tertiary amine derivatives **7** and **8**, as well as urea analogue **9**, showed substantially reduced activity (Table 1).

**Table 1.**
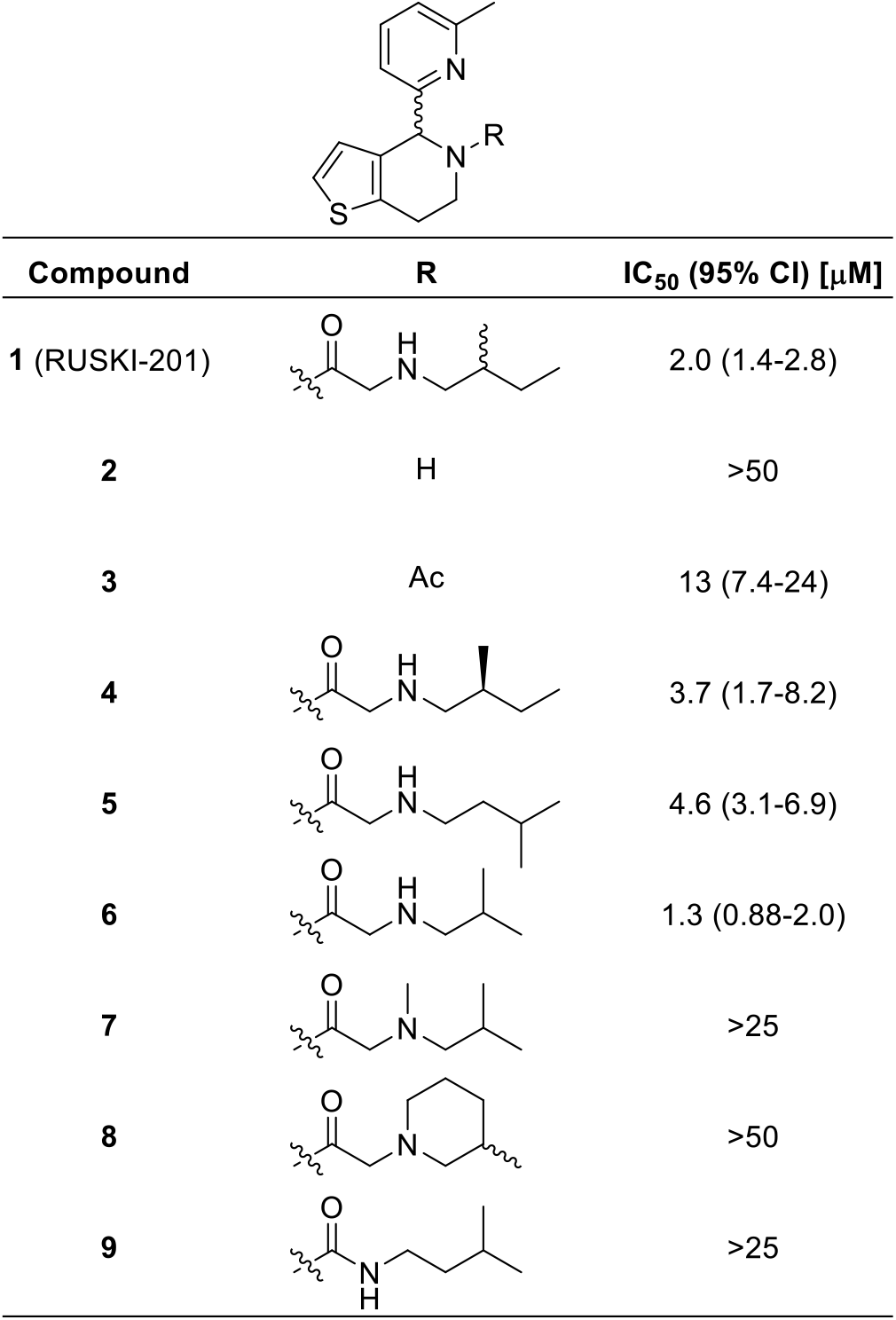
Structure-activity relationship investigation of the amide substituent of **1** (RUSKI-201). Data represent mean and 95% confidence interval (CI, n = 3).

|  |  |  |
| --- | --- | --- |
| Compound | R | IC<sub>50</sub> (95% CI) [μM] |
| 1 (RUSKI-201) |  | 2.0 (1.4-2.8) |
| 2 | H | >50 |
| 3 | Ac | 13 (7.4-24) |
| 4 |  | 3.7 (1.7-8.2) |
| 5 |  | 4.6 (3.1-6.9) |
| 6 |  | 1.3 (0.88-2.0) |
| 7 |  | >25 |
| 8 |  | >50 |
| 9 |  | >25 |

The most potent compound, **6**, contains an undefined stereocentre in the dihydrothieno[3,2-*c*]pyridine core, which raised the possibility that only one enantiomer may fit the binding site in HHAT. **(+/−)−6** was therefore purified by chiral preparative HPLC to obtain **(+)−6** and **(−)−6** in >99:1 enantiomeric ratio (Figure S1). **(−)−6** displayed no HHAT inhibition, whilst **(+)−6** displayed two-fold increased potency compared to **(+/−)−6** (Table 2). The lead inhibitor **(+)−6**, which we here term **IMP-1575**, is the highest potency small-molecule HHAT inhibitor to-date (IC_50_ = 0.75 μM, 95% CI = 0.49-1.1 μM). **IMP-1575** is an oil, precluding determination of absolute stereochemistry by X-ray methods despite extensive efforts. Subsequent studies will be required to confirm the stereochemistry of this molecule.

**Table 2.**
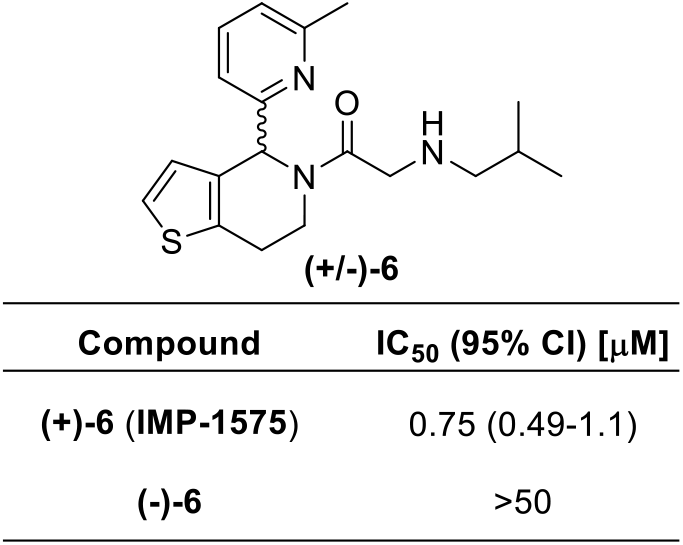
Development of single-enantiomer HHAT inhibitor **(+)−6** (**IMP-1575**). **(−)−6** does not inhibit HHAT, whereas **(+)−6** (**IMP-1575**) is twice as potent as **(+/−)−6**. Data represent mean and 95% CI (n = 3).

|  |  |
| --- | --- |
| Compound | IC<sub>50</sub> (95% CI) [μM] |
| (+)-**6** (**IMP-1575**) | 0.75 (0.49-1.1) |
| (-)-**6** | >50 |

Collectively, SAR data demonstrated the importance of the secondary amine in HHAT binding, and tolerance for alteration of the alkyl chain. These new insights led to design of photocrosslinking chemical probe **10** to investigate the small-molecule binding site in HHAT (Figure 2A). In probe **10** the secondary amine moiety was exchanged for a diazirine group, and the terminal alkyl chain replaced with an alkyne. The diazirine allows photoactivated chemical crosslinking to nearby residues in HHAT, with the alkyne allowing biorthogonal functionalization via copper^I^-catalyzed azide-alkyne cycloaddition (CuAAC, ‘click chemistry’) for analysis. The sidechain was synthesized (Scheme S3) and coupled to **2** (Scheme S2). Pleasingly, probe **10** retained inhibitory activity against HHAT in the Acyl-cLIP assay (IC_50_ = 2.4 μM, 95% CI = 1.8-3.3 μM, Figure 2B) demonstrating the key functional groups were well tolerated. Photoactivation of **10** by UV irradiation (365 nm) was monitored by LC/MS, demonstrating an activation half-life of 32 s (Figure S2). Protein crosslinking by **10** was next investigated through CuAAC functionalization with a fluorophore and in-gel fluorescence readout. As an integral membrane protein, HHAT presents substantial challenges in sample handling. Purified HHAT migrates as a diffuse band in SDS-PAGE,^[14]^ and therefore crosslinking was performed in lysate from HEK293a cells overexpressing HHAT-FLAG-His_8_ (HEK293a HHAT^+^) as the protein migrates as a well-defined, predominantly monomeric species.^[15]^ Further, HHAT cannot be precipitated to remove excess fluorophore which otherwise binds non-specifically (data not shown). Fluorogenic azide CalFluor-647 (Figure S3)^[16]^ was therefore used to reduce background fluorescence from dye that had not reacted with alkyne probe. **10** (10 and 30 μM) was incubated in HEK293a HHAT^+^ lysate with 5 min UV irradiation. Functionalization with CalFluor-647 by CuAAC followed by in-gel fluorescence analysis indicated crosslinking predominantly to a species at ^~^50 kDa, consistent with the apparent molecular weight of HHAT as assessed by anti-FLAG immunoblotting (Figure 2C).

**Figure 2.**
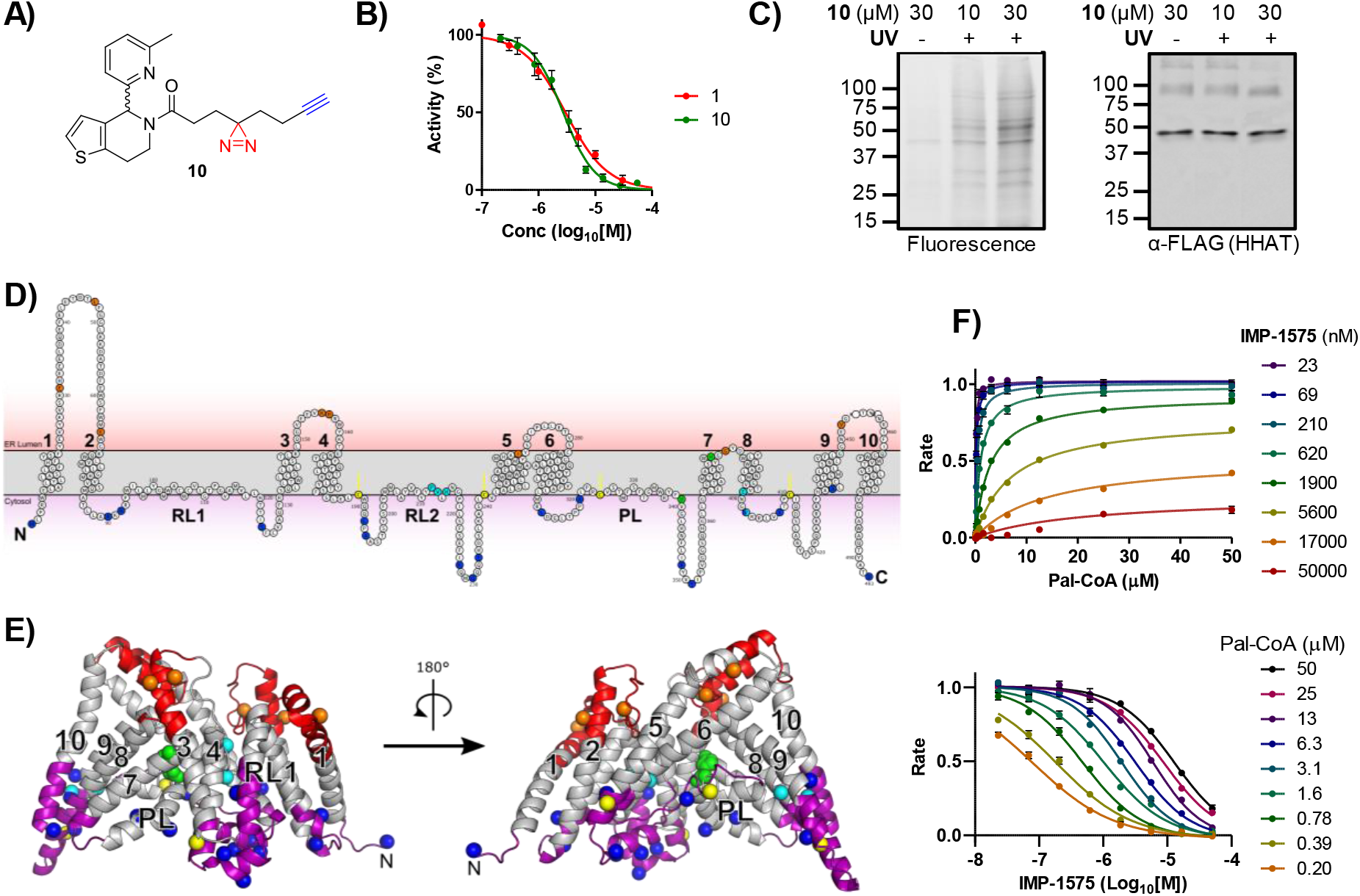
Photochemical probe identification of binding site in HHAT. A) Structure of photocrosslinking probe **10**. B) Acyl-cLIP assays showing equal potency between **1** (RUSKI-201) and photochemical probe **10**. Data represent mean and SEM (n = 3). C) UV-crosslinking (365 nm, 5 min) of **10** to HEK293a cell lysate expressing HHAT-FLAG-His_8_. Labelled proteins were functionalised with CalFluor647 by copper^I^-catalysed azide-alkyne cycloaddition. Analysis by in-gel fluorescence and α-FLAG immunoblotting showed crosslinking predominantly to a band at a similar molecular weight to monomeric HHAT. D) Topology model of HHAT showing cytosolic (purple) and ER lumen (red) loops. Key residues are coloured as catalytic His379 and Asp339 (green), probe-modified Pro212, Val213, His215, Glu399 and Val402 (cyan), palmitoylated cysteines (yellow), and residues that have had their topology experimentally determined as cytosolic (blue) or luminal (orange). E) Homology model of HHAT, coloured as in (D), showing probe modified residues Pro212, Val213, His215 in proximity to the central catalytic site on the cytosolic face of HHAT. F) Effect of **IMP-1575** on HHAT kinetics, showing increased inhibitor concentration caused increased Pal-CoA *K*_M_ (top), and increased Pal-CoA concentration decreased **IMP-1575** potency (bottom). Analysis with a mixed model equation for inhibition generated α = 470 μM (95% CI = 320-620 μM), demonstrating competitive inhibition (*K*_i_ = 38 nM, 95% CI = 29-46 nM). Data represent mean and SEM (n = 3)

Having identified suitable conditions for photocrosslinking, identification of the HHAT small-molecule binding site was sought through crosslinking with protease digestion and tandem LC-MS/MS analysis. To achieve maximum sequence coverage purified HHAT (1 μg)^[14]^ was digested with trypsin, Glu-C, Proteinase K, chymotrypsin, or combinations of Lys-C and trypsin, or ProteaseMAX and trypsin. Filter-aided sample preparation (FASP)^[17]^ was used to remove the *n*-dodecyl-β-D-maltopyranoside (DDM) detergent, and reduce and alkylate peptides prior to analysis. Moderate-to-low sequence coverage was achieved with all proteases (Table S1), with chymotrypsin having greatest coverage (51%). Combined, chymotrypsin, Proteinase K (41%), and trypsin (35%) gave 68% overall sequence coverage (Figure S4). Purified HHAT and **10** (25 μM) were irradiated for 3 min, followed by FASP processing and digestion with trypsin, chymotrypsin, or Proteinase K. LC-MS/MS identified two peptides with increased mass consistent with crosslinking to **10**, and MS/MS fragmentation indicated modification of residues Pro212, Val213, His215, Glu399 and Val402 (Figure S5).

To investigate how the identified binding residues may be involved in HHAT’s structure and function, residue positions were compared to the experimentally determined membrane topology of HHAT (Figure 2D).^[9]^ Probe-binding residues were located in re-entrant loop 2 (Pro212, Val213 and His215), which is positioned near the cytosolic face of the membrane, or immediately following transmembrane helix 8 (Glu399 and Val402), which is also positioned on the cytosolic membrane face. The binding residues were distant in the HHAT sequence from the signature MBOAT catalytic residues (Asp339 and His379), therefore a structural model for HHAT was generated based on sequence homology to DltB^[5]^ (Figure 2E). Phyre2, SwissModel and Robetta (old and new) servers were used to generate homology models of HHAT. Models were initially screened based on agreement with previous topology data.^[9]^ Interestingly, all models shared the same fold between residues 94-206, with greater variability in the final two predicted transmembrane helices at the N- and C-termini. A final model was selected based on optimal hydrophobic packing of the terminal membrane-spanning helices (Figure 2E). Predicted transmembrane regions from topology studies were arranged around a central pore (Figure 2E, shown in grey). Signature MBOAT His379 was located at the center of the pore in close proximity to the essential Asp339, suggesting a central catalytic site. Inhibitor-binding residues Pro212, Val213, and His215 were located adjacent to the proposed catalytic site, whereas residues Glu399 and Val402 were not located in proximity to the proposed catalytic site in this model, likely reflecting lower accuracy of the model in this region (Figure 2E). Collectively, this analysis suggested that the HHAT inhibitor-binding site may be located on the cytosolic side of the ER membrane in proximity to the catalytic site. As Pal-CoA is expected to approach HHAT from the cytosolic side of the membrane, **IMP-1575** inhibition (50-0.023 μM) was analyzed for effects on HHAT kinetics at varying Pal-CoA concentrations (50-0.19 μM, Figure 2F). Analysis using a mixed model equation for inhibition^[18]^ generated α = 470 (95% CI = 320-620), indicative of a competitive mode of inhibition for **IMP-1575** with respect to Pal-CoA (*K*_i_ = 38 nM, 95% CI = 29-46 nM). Taken together, these data support identification of a small-molecule binding site on the cytosolic face of HHAT that competes with Pal-CoA binding to disrupt enzymatic function with high affinity.

We present here a new paradigm in rational development of photocrosslinking chemical probes for MBOATs that can identify small-molecule binding sites on these very challenging but important targets. Probe **10** was designed based on new SAR insights into known inhibitor **1**, which retained required HHAT inhibitory activity (Figure 2). Probe **10** was used here to identify a small-molecule inhibitor binding site in an MBOAT for the first time. Other mammalian MBOATs, such as PORCN and GOAT, have known inhibitors and their investigation may be expedited by technical advancements reported here, including fluorogenic click regents for SDS-PAGE analysis, and FASP processing in combination with multiple proteases to increase sequence coverage. Photochemical tools based on substrates may further allow mapping of key functional sites in HHAT and other MBOATs. The limitations of biochemical topology analysis and homology models for inhibitor design are significant, therefore structural information for HHAT and other MBOATs such as PORCN and GOAT remains an important goal for inhibitor development. We further provide **IMP-1575**, a single enantiomer small molecule, as the most potent HHAT inhibitor reported to date that competes with Pal-CoA (*K*_i_ = 38 nM, Figure 2). Previous studies of this chemical series have suggested compounds are noncompetitive with Pal-CoA (RUSKI-43, *K*_i_ = 6.9 μM);^[19]^ however, these experiments did not test inhibitor concentrations below 10 μM, therefore preventing accurate mechanistic determination. Future studies of HHAT enzymology could take advantage of **IMP-1575**, with the inactive enantiomer serving as an ideal control. In this regard, the improved synthetic routes to HHAT inhibitors (Schemes S1 and S2) presented here will significantly accelerate future development of this series. In summary, we present chemical tools and methodology to provide insight into HHAT which may expedite future studies and drug discovery efforts against this important target class.

## Acknowledgements

This study was supported by Cancer Research UK grants (C6433/A16402 to A.I.M. and E.W.T; C29637/A20183 to E.W.T., and C20724/A14414 and C20724/A26752 to C.S.), the European Union Framework Program 7 (Marie Curie Intra European Fellowship to M.R. and E.W.T., and to M.M. and E.W.T.), the European Research Council under the European Union’s Horizon 2020 research and innovation programme (647278 to C.S.), the Wellcome Trust (DPhil studentships 102749/Z/13/Z to L.S. and BST00110.B500.04 to C.E.C.). TLH is a Career Development Fellow funded by the Department of Pharmacology, University of Oxford.

## Notes

### Competing Interest Statement

The authors have declared no competing interest.

