## Supporting Information for "Photochemical probe identification of the small-molecule binding site in a mammalian membrane-bound *O*-acyltransferase"

**Supplementary Data**

Supplementary Figure S1. Chiral HPLC traces for separation of (+)-6 (IMP1575) and (-)-6. . 6

### Supplementary Schemes

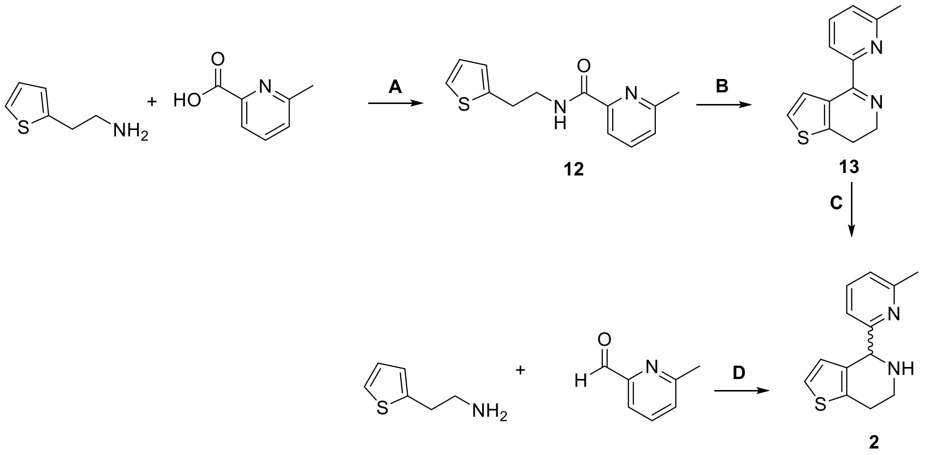

| Route | 2 yield (%) |
|---|---|
| A, B, C | 9 |
| D | 54 |

**Supplementary Scheme S1. Synthesis of key intermediate 2.**

General procedures: **A)** PyBOP, DIPEA, DMF, room temperature, overnight; **B)** $POCl_3$, $P_2O_5$, toluene, microwave (140 °C); **C)** $NaBH_4$, room temperature, 1 h;[1] **D)** EtOH, TEA, room temperature, 15 h, then TFA, room temperature, 30 min. The new route for synthesis of **2** via general procedure D is both quicker and higher yielding.

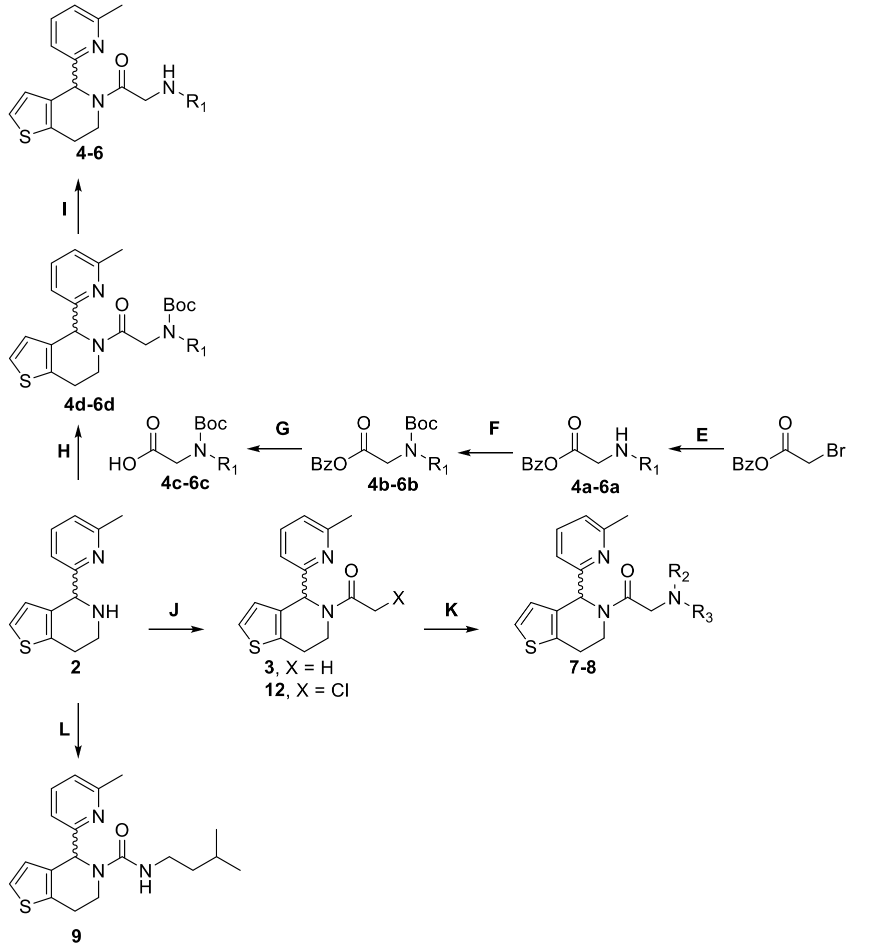

**Supplementary Scheme S2. Synthesis of analogues 3-9.**

General procedures: **E)** room temperature, overnight; **F)** amine, TEA, $Boc_2O$, 0 °C, 15 min; **G)** 10% Pd/C, $NH_4HCO_3$, 75 °C, 30 min; **H)** EDC, HOBt, DIPEA; **I)** TFA, room temperature, 3 h; **J)** acyl chloride, THF, DIPEA, 0 °C to room temperature, 1 h; **K)** amine, $K_2CO_3$, MeCN, 60 °C, overnight; **L)** CDI, room temperature, 1 h.

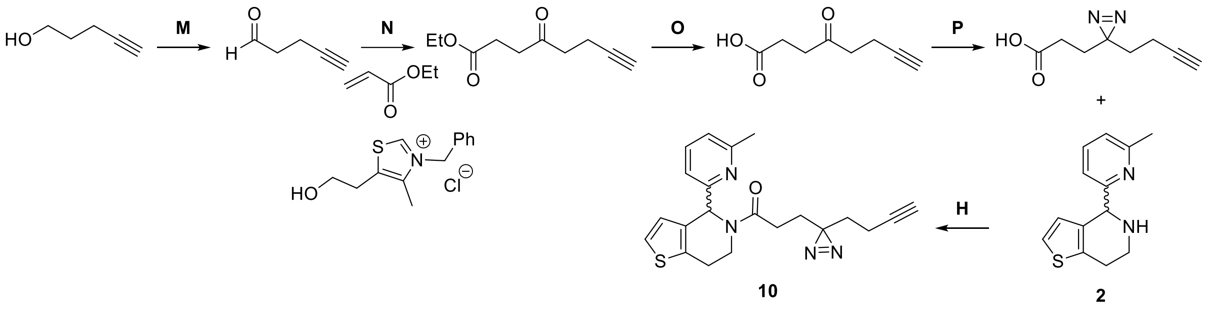

**Supplementary Scheme S3. Synthesis of probe 10.**

General procedures: **M)** Dess-Martin periodinane, dichloromethane, 0 °C to room temperature, 2 h; **N**) TEA, 1,4-dioxane, 84 °C, 54 h; **O**) LiOH, $H_2O$, MeOH, room temperature, overnight; **P**) 7 M $NH_3$, MeOH, 0 °C, 3 h, then $NH_2OSO_3$, MeOH, 0 °C to room temperature, overnight, then $I_2$, DIPEA, MeOH, 0 °C, 2 h.[2]

### Supplementary Figures

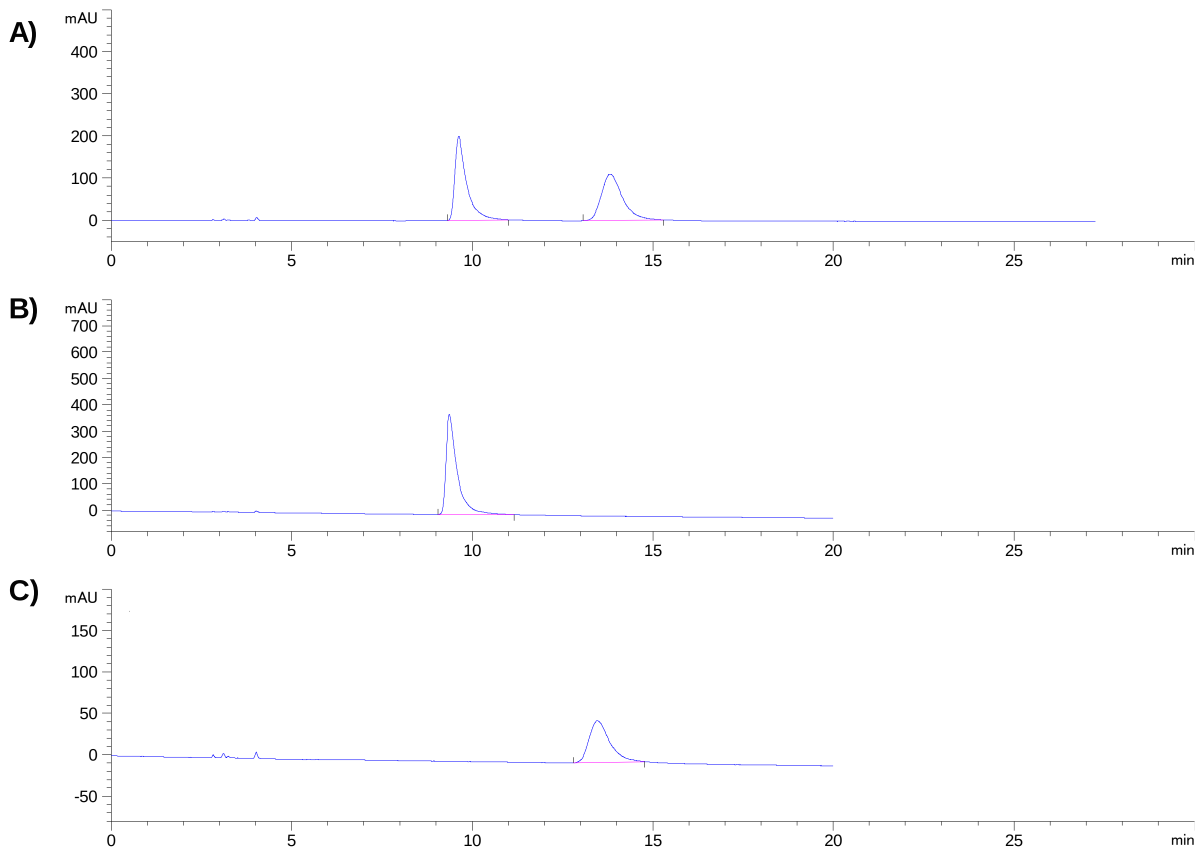

| Chromatogram | Compound | Peak (min) | Area (%) |
|---|---|---|---|
| **A** | **(+/-)-6** | 9.6 | 50 |
| | | 13.8 | 50 |
| **B** | **(+)-6 (IMP-1575)** | 9.4 | 100 |
| **C** | **(-)-6** | 13.5 | 100 |

**Supplementary Figure S1. Chiral HPLC traces for separation of (+)-6 (IMP1575) and (-)-6.**

A) Analytical chiral HPLC trace for **(+/-)-6**. B) Analytical chiral HPLC trace for **(+)-6** (**IMP-1575**) following preparative chiral HPLC. C) Analytical chiral HPLC trace for **(-)-6** following preparative chiral HPLC.

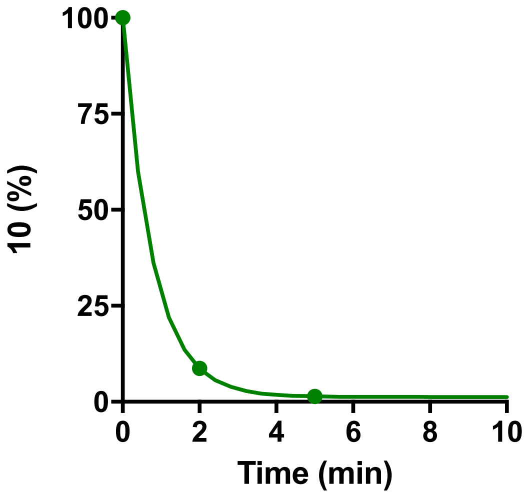

**Supplementary Figure S2. Rate of photoactivation of probe 10.**

Probe **10** was UV irradiated (365 nm) and the presence of the starting material monitored by UV peak area by LC-MS. Data expressed as percentage of **10** remaining.

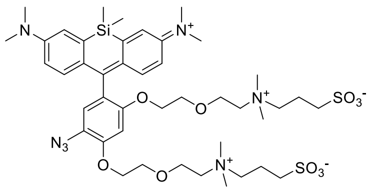

**Supplementary Figure S3. Structure of CalFluor647.**

Structure of CalFluor647 fluorogenic dye which undergoes a turn-on in fluorescence after copper$^{I}$-catalysed azide-alkyne cycloaddition (CuAAC).[3]

1 MLPRWELALY LLASLGFHFY SFYEVYKVSR EHEEELDQEF ELETDTLFGG LKKDATDFEW SFWMEWGKQW LVWLLLGHMV

21 VSQMATLLAR KHRPWILMLY GMWACWCVLG TPGVAMVLLH TTISFCVAQF RSQLLTWLCS LLLLSTLRLQ GVEEVKRRWY

161 KTENEYYLLQ FTLTVRCLYY TSFSLELCWQ QLPAASTSYS FPWMLAYVFY YPVLHNGPIL SFSEFIKQMQ QQEHDSLKAS

241 LCVLALGLGR LLCWWWLAEL MAHLMYMHAI YSSIPLLETV SCWTLGGLAL AQVLFFYVKY LVLFGVPALL MRLDGLTPPA

321 LPRCVSTMFS FTGMWRYFDV GLHNFLIRYV YIPVGGSQHG LLGTLFSTAM TFAFVSYWHG GYDYLWCWAA LNWLGVTVEN

401 GVRRLVETPC IQDSLARYFS PQARRRFHAA LASCSTSMLI LSNLVFLGGN EVGKTYWNRI FIQGWPWVTL SVLGFLYCYS

481 HVGIAWAQTY ATD

**Supplementary Figure S4. Coverage of HHAT sequence.**

Sequence coverage of HHAT after protease digestion. Underlined sequence represents detection of a peptide after digestion with trypsin (purple, 35%), chymotrypsin (green, 51%) or proteinase K (red, 41%). Collectively the three proteases allow detection of 68% of the amino acid sequence of HHAT. Signature MBOAT residues His379 and Asp339 are shown in green, with probe-modified residues Pro212, Val213, His215, Glu399 and Val402 shown in blue.

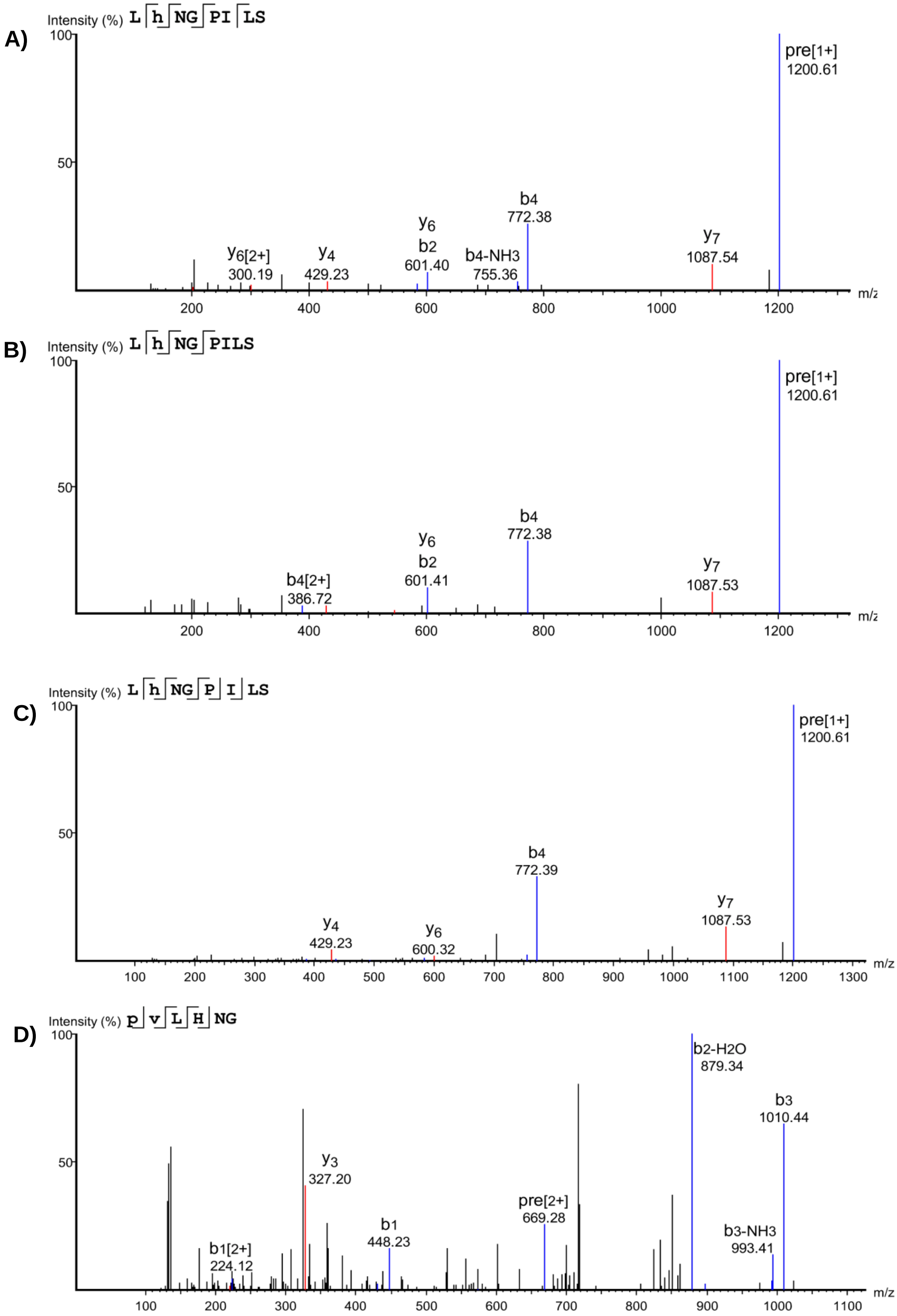
A)
Intensity (%) L h NG PI LS
100
50
y6[2+]
300.19
y4
429.23
y6
b2
601.40
b4-NH3
755.36
b4
772.38
y7
1087.54
pre[1+]
1200.61
200
400
600
800
1000
1200
m/z
B)
Intensity (%) L h NG PILS
100
50
b4[2+]
386.72
y6
b2
601.41
b4
772.38
y7
1087.53
pre[1+]
1200.61
200
400
600
800
1000
1200
m/z
C)
Intensity (%) L h NG P I LS
100
50
y4
429.23
y6
600.32
b4
772.39
y7
1087.53
pre[1+]
1200.61
100
200
300
400
500
600
700
800
900
1000
1100
1200
1300
m/z
D)
Intensity (%) p v L H NG
100
50
b1[2+]
224.12
y3
327.20
b1
448.23
pre[2+]
669.28
b2-H2O
879.34
b3-NH3
993.41
b3
1010.44
100
200
300
400
500
600
700
800
900
1000
1100
m/z

E)

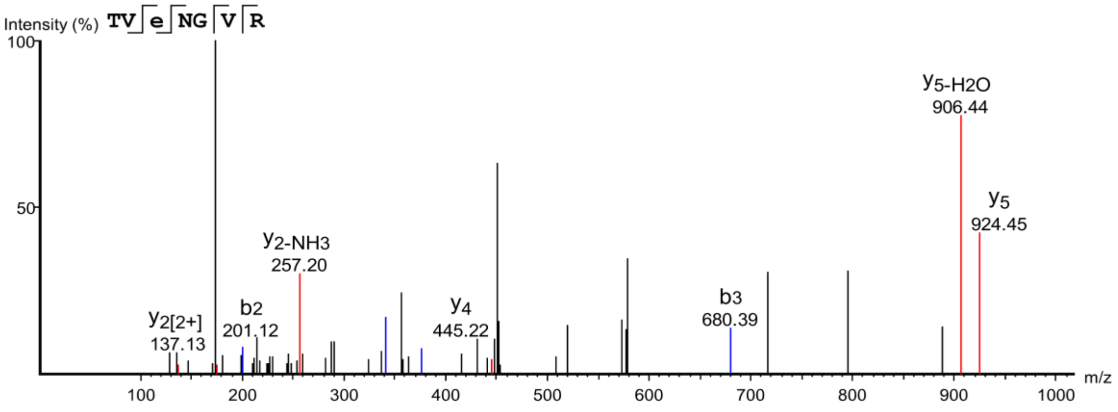

F)

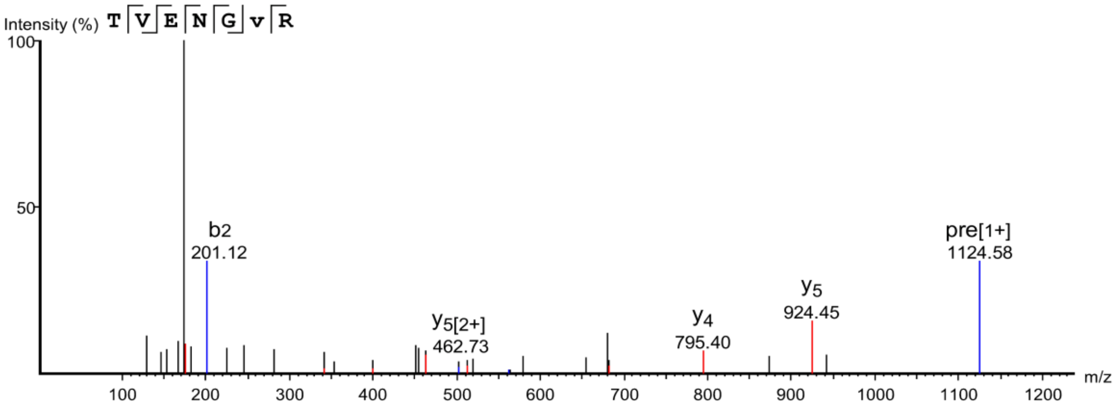

**Supplementary Figure S5. LC-MS/MS detection of peptides modified by probe 10.**

MS/MS fragmentation of peptides with molecular weight increases consistent with crosslinking to **10**. A-C) LhNGPILS for His215 modification. D) pvLHNG for Pro212, Val213 modification. E) TVeNGVR for Glu399 modification. F) TVENGvR for Val402 modification.

### Supplementary Tables

| Protease(s) | Coverage (%) |
|---|---|
| Trypsin | 35 |
| Glu-C | 5 |
| Proteinase K | 41 |
| Chymotrypsin | 51 |
| Lys-C and trypsin | 36 |
| ProteaseMAX and trypsin | 35 |

**Table S1. HHAT sequence coverage following protease digestion.**

Sequence coverage following digestion of HHAT (1 µg) with either trypsin, Glu-C, Proteinase K, chymotrypsin, or combinations of Lys-C and trypsin, and ProteaseMAX and trypsin. Data represent percentage of HHAT sequence with a statistical significance of -LogP = 20

### Materials and Methods

#### Abbreviations

DDM (*n*-dodecyl-β-D-maltopyranoside); DIPEA (*N*,*N*-diisopropylethylamine); DMF (dimethylformamide); DTT (dithiothreitol); EDTA (ethylenediaminetetraacetic acid); FAM (5/6-carboxyfluorescein); HATU (1-[bis(dimethylamino)methylene]-1*H*-1,2,3-triazolo[4,5-b]pyridinium 3-oxid hexafluorophosphate); HBTU (*N*,*N*,*N*′,*N*′-tetramethyl-*O*-(1*H*-benzotriazol-1-yl)uronium hexafluorophosphate); HCTU (1-[bis(dimethylamino)methylen]-5-chlorobenzotriazolium 3-oxide hexafluorophosphate); HEPES (4-(2-hydroxyethyl)-1-piperazineethanesulfonic acid); HOBt (hydroxybenzotriazole; MMT (monomethoxytrityl); MTS (3-(4,5-dimethylthiazol-2-yl)-5-(3-carboxymethoxyphenyl)-2-(4-sulfophenyl)-2H-tetrazolium); NMM (*N*-methylmorpholine); NMP (*N*-methyl-2-pyrrolidone); PBS (phosphate buffered saline); PBS-T (phosphate buffered saline plus Tween-20 (0.1%)); RT (room temperature); TBTA (tris[(1-benzyl-1H-1,2,3-triazol-4-yl)methyl]amine); TCEP (tris(2-carboxyethyl)phosphine hydrochloride);TFA (trifluoroacetic acid); TFE (trifluoroethanol); TIS (triisopropylsilane); TMP (trimethoxyphenylthio); TRIS (tris(hydroxymethyl)aminomethane).

#### General Information

All reagents were purchased from commercial sources (Sigma-Aldrich, Fisher Scientific, Acros Organics, Novabiochem) and used without further purification.

Pal-CoA (Sigma Aldrich) was stored in NaOAc (0.1 M, pH 6.0) at -80 °C. RUSKI-201 (**1**) was synthesised as previously described.[1b] The peptide comprising of residues 24-43 of SHH (residues 1-10 of the mature protein) labelled with 5/6-carboxyfluorescein (SHH-FAM) was prepared as previously described.[4]

Mixtures described as % represent v/v unless otherwise stated.

#### Biological and Biochemical Methods

#### HHAT Purification

Full-length human HHAT (UniProt Q5VTY9) C-terminally fused with a 1D4 epitope tag was expressed and purified via antibody purification with Rho-1D4 antibody (University of British Columbia) coupled to CNBr-activated sepharose beads (GE Healthcare) as previously described.[5] Purified HHAT 0.1-0.2 mg/mL in storage buffer (10 mM HEPES, 150 mM NaCl, 5% glycerol, 0.025% (w/v) DDM, 0.0006% (w/v) cholesterol-hemisuccinate, pH 7.5) was flash-frozen in liquid nitrogen and stored at -80 °C.

#### Acyl-cLIP Assay

The Acylation-coupled Lipophilic Induction of Polarisation (Acyl-cLIP) was performed as previously described.[5] Briefly, HHAT (450 µg/mL) in solubilisation buffer (20 mM HEPES, 350 mM NaCl, 1% (w/v) DDM, 5% glycerol, pH 7.3) was diluted 1:10 in SHH-FAM (2 µM) in reaction buffer (100 mM MES, 20 mM NaCl, 1 mM DTT, 1 mM TCEP, 0.1% (w/v) BSA, pH 6.5). The HHAT/SHH-FAM solution (12 µL/well) was dispensed into a black 384-well plate (Corning,

3575). The reaction was initiated by addition of Pal-CoA (7.5 µM) in reaction buffer (8 µL/well), and fluorescence anisotropy measured every 1 min for 1 h on an EnVision Xcite 2104 (PerkinElmer). Measurement used a FITC FP D505fp mirror module, FITC FP 480 nm excitation filter (30 nm bandwidth), FITC FP P-pol 535 nm first emission filter (40 nm bandwidth), and FITC FP S-pol 535 nm second emission filter (40 nm bandwidth), measurement height = 9.52 mm, 8 flashes/well, PMT gain = 319, and G-factor = 1.09. Fluorescence anisotropy was calculated using equation 1:

$$(1)\ FA = \frac{S - GP}{S + 2GP}$$

Where S is signal in the S-channel, P is signal in the P-channel, and G is the G-factor.

Assay data were evaluated using Microsoft Excel 2013 and GraphPad Prism 5.0. Initial rate constants were determined by linear regression, normalized to DMSO negative controls and background corrected to positive controls with solubilization buffer instead of HHAT. The half-maximal inhibitory concentration ($IC_{50}$) was calculated using a 4-parameter sigmoidal dose-response curve fit. Determination of the mechanism of inhibition of IMP-1575 relative to Pal-CoA was performed by fitting equation 2 for a mixed model of inhibition[6] using nonlinear regression:

$$(2)\ v = \frac{\dfrac{V_{max}}{\left(1 + \dfrac{[I]}{\alpha K_i}\right)}[S]}{[S] + K_M\left(\dfrac{\left(1 + \dfrac{[I]}{K_i}\right)}{\left(1 + \dfrac{[I]}{\alpha K_i}\right)}\right)}$$

Where v is the observed rate of reaction, $V_{max}$ is the maximum rate, [I] is the inhibitor concentration, [S] is the substrate concentration, $K_m$ is the substrate concentration at ½ $V_{max}$, $K_i$ is the affinity of the inhibitor for the free enzyme, and α is the ratio of the affinity of the inhibitor for the enzyme-substrate complex ($K_i$') to $K_i$ (typically noted as $K_i$'/$K_i$).

#### Photoactivation Analysis by LC/MS

A solution of **10** (100 µM) in $H_2O$ was cooled on ice and irradiated using a custom-made UV light source (365 nm, 25 mW/cm$^2$) for 0, 2 and 5 min. LC-MS analysis was performed on a Waters preparative HPLC-MS equipped with a Waters XBridge C18 4.6 × 100 mm column with 1.2 mL/min flow rate, 5-98% MeOH in $H_2O$ gradient over 18 min. All eluents were supplemented with 0.1% formic acid.

#### Click Chemistry and SDS-PAGE Analysis

HEK293a cells overexpressing HHAT-FLAG-His (HEK293a-HHAT$^+$) were cultured as previously described.[9] Cells were resuspended in solubilisation buffer (300 µL) and lysed via sonication with an ultrasonic homogenizer (20 kHz, 10 ×5 s pulses). After centrifugation (5 min, 16,200 g, 4 °C), the soluble lysate was removed and diluted with solubilisation buffer to

1 mg/mL. Probe **10** (1 or 3 mM, 1 µL) or DMSO (1 µL) was added to lysate (99 µL) and samples UV irradiated as previously described. 'Click-mix' was prepared by adding $CuSO_4$ (50 mM in $H_2O$, 2 vol) to CalFluor-647 (10 mM in DMSO, 1 vol), followed by TCEP (50 mM in water, 2 vol) and TBTA (10 mM in DMSO, 1 vol). The 'Click-mix' was vortexed (2 min, room temperature), then added to lysates (5 µL/sample), and samples vortexed (1 h, room temperature) whilst covered to exclude light. The reaction was quenched by addition of EDTA (10 mM final concentration) and vortexed (5 min, room temperature). Samples were supplemented with SDS-PAGE loading buffer (5 × final concentration stock: 20% β-mercaptoethanol, 0.02% (w/v) bromophenol blue, 30% glycerol, 10% (w/v) SDS, 250 mM Tris, pH 6.8). Following electrophoresis, CalFluor647 in-gel fluorescence was recorded using a Typhoon imager (GE Healthcare). Following imaging, proteins were transferred to a nitrocellulose membrane and probed by α-FLAG immunoblotting as previously described.[4]

#### Protease Digestion and Sequencing

HHAT (10 µL, 0.1 mg/mL) was diluted 1:10 in proteomics sample buffer (100 mM MES, 20 mM NaCl, 1 mM TCEP, 1 mM DTT, 0.1% (w/v) DDM, pH 6.5), transferred to a 10 kDa molecular weight cut-off (MWCO) Vivaspin 500 concentrator (Sartorius) and incubated for 15 min on ice. Urea solution (200 µL, 8 M urea in 100 mM Tris, pH 8.5, kept on ice) was added and the samples were centrifuged (14,000 g, 10 min, 4 °C). Iodoacetamide (500 mM, kept on ice) was added to a final concentration of 50 mM and the mixture was vortexed for 1 min before incubation in the dark for 20 min. The concentrators were washed with urea solution (3 × 100 µL) and with AMBIC solution (50 mM $NH_4HCO_3$, pH 8.5, 3 × 100 µL) via centrifugation (10,000 g, 3 min, 4 °C).

Digestions were performed using (1) sequencing grade modified trypsin (Promega) (1:10, 0.5 µL of 0.2 µg/µL stock); (2) Glu-C (Promega) (1:20, 0.5 µL of 0.1 µg/µL stock); (3) Proteinase K (New England Biolabs) (1:20, 2.5 µL of 20 ng/µL stock); (4) chymotrypsin (Promega) (1:20, 0.5 µL of 0.1 µg/µL stock); (5) Lys-C (1:10, 0.5 µL of 0.2 µg/µL stock) and trypsin (1:10, 0.5 µL of 0.2 µg/µL stock); (6) Protease MAX (1:10, 1 µL of 0.1 µg/µL stock) and trypsin (1:10, 0.5 µL of 0.2 µg/µL stock). All samples were shaken overnight at 37 °C except for chymotrypsin (25 °C, overnight). Digested peptides were collected into a clean Vivacon tube with 40 µL washes of AMBIC solution (50 mM) and NaCl (0.5 M).

Samples were acidified with 0.5% TFA and loaded onto stage tips containing three SDB-XC poly(styrenedivinyl-benzene) copolymer discs (Merck). The stage tipping procedure was performed as described previously.[7] Peptides were eluted in 80% acetonitrile in $H_2O$ and the solvent removed using a Savant SPD1010 SpeedVac® Concentrator at 45 °C.

Peptide samples were dissolved in LC-MS grade $H_2O$ containing 2% acetonitrile and 0.5% TFA (15 µL) for nano-LC-MS/MS

#### Crosslinking for Binding Site Identification

HHAT (100 µL, 0.01 mg/mL) in proteomics sample buffer was incubated with photochemical probe **10** (25 µM), on ice for 15 min in 10 kDa MWCO Vivaspin 500 concentrator (Sartorius).

The samples were irradiated for 3 min as previously described. Samples were prepared for nanoLC-MS/MS analysis as previously described.

**NanoLC-MS/MS Data Acquisition**

Peptide samples (3 µL) were separated on an EASY-Spray™ Acclaim PepMap C18 column (50 cm × 75 µm inner diameter, 3 µm particle size, Thermo Fisher Scientific) using a 2 h linear gradient separation of 0-100% solvent B (80% acetonitrile supplemented with 0.1% formic acid): solvent A (2% acetonitrile supplemented with 0.1% formic acid) at a flow rate of 250 nL/min. The liquid chromatograph was coupled to a Q Exactive mass spectrometer via an easy-spray ion source which was operated in data-dependent mode with survey scans acquired at a resolution of 70,000 at m/z = 200. Scans were acquired from 350 to 1650 m/z. Up to 10 of the most abundant isotope patterns with charge +2 or higher from the survey scan were selected with an isolation window of 1.6 m/z and fragmented by higher-energy collisional dissociation with normalized collision energy of 25. The maximum ion injection times for the survey scan and the MS/MS scans (acquired with a resolution of 35,000 at m/z = 200) were 20 and 120 ms, respectively. The ion target value for MS was set to $10^6$ and for MS/MS to $10^5$, and the intensity threshold was set to $8.3 \times 10^2$.

**Peptide Search and Data Analysis**

Raw files were analysed in PEAKSStudio8.5 and searched against the HHAT sequence. Cysteine carbamidomethylation, methionine oxidation and N-terminus acetylation were selected as variable modifications. Probe modification was set as fixed modification for crosslinking experiments. Up to two missed cleavages were allowed. –10logP value was set to ≥20 for peptides Other parameters were used as preset in the software (precursor mass error tolerance 15 ppm, fragment mass error tolerance 0.5 Da, precursor mass type monoisotopic, maximum variable PTM per peptide 3, maximum missed cleavage 2, and non-specific cleavage 1).

**HHAT Modelling**

A set of homology models for human HHAT (Uniprot ID: Q5VTY9; Genbank ID: NM_001122834.3, natural variant VAR_024743) were built using the Robetta Web Server (http://old.robetta.org/) based on the structure of the bacterial MBOAT member DltB (PDB ID: 6BUH).[8] A final model was selected based on agreement with topological data.[9]

**Chemical Synthesis**

**General Information**

Analytical chiral HPLC was performed on an Agilent 1260 Infinity Series equipped with a CHIRALPAK-IA 4.6 mm × 250 mm (eluent: hexane:propan-2-ol 90:10 or 80:20; flow rate 1 mL/min, Method A), a CHIRALPAK-ID 4.6 mm × 250 mm (eluent: isocratic hexane:propan-2-ol 90:10; flow rate 1 mL/min, Method B) or a CHIRALPAK-IF 4.6 mm × 250 mm (eluent: isocratic hexane:propan-2-ol 90:10 or 80:20; flow rate 1 mL/min, Method C). Preparative chiral HPLC was performed on an Agilent 1200 Series equipped with a CHIRALPAK-IF

250 mm × 20 mm column (eluent: isocratic hexane:propan-2-ol 90:10, flow rate 18 mL/min, Method D; isocratic hexane:propan-2-ol 80:20, flow rate 18 mL/min, Method E). Optical rotations were measured using a BS ADP440+ polarimeter.

**Synthetic procedures**

**Procedures A-C (synthesis of 2)**

General procedures A-C were performed as previously described.[1]

**Procedure D (synthesis of 2)**

To a stirred solution of 6-methylpicolinaldehyde (1 eq) in ethanol (3.5 mL) was added 2-(thiophen-2-yl)ethan-1-amine (1 eq) and triethylamine (0.2 mL) and the solution stirred for 15 h. The solvent was removed under reduced pressure and the residue was dissolved in TFA (15 mL) and stirred at room temperature for 30 min. The solvent was removed under reduced pressure and the residue was dissolved in dichloromethane, washed with 2 N NaOH (aq), and the aqueous layer extracted with dichloromethane. The organics were combined, dried over $MgSO_4$, filtered and concentrated under reduced pressure to afford a brown oil which was recrystallised from dichloromethane/hexane.

**Procedure E (side chain synthesis, first step)**

Benzyl bromoacetate (1 eq) was added dropwise to the corresponding primary amine (3 eq) in dry tetrahydrofuran (THF) at 0 °C under nitrogen atmosphere. The solution was stirred overnight at room temperature. The solvent was removed under reduced pressure and the residue purified by column chromatography.

**Procedure F (side chain synthesis, Boc protection)**

Triethylamine (1.5 eq) was added to the secondary amine (1 eq) dissolved in dry methanol at 0 °C. Subsequently, di-*tert*-butyl dicarbonate (1 eq) dissolved in dry methanol was added and the mixture was stirred under a nitrogen atmosphere at 0 °C for 15 min. The solvent was removed under reduced pressure and the residue purified by column chromatography.

**Procedure G (side chain synthesis, benzyl deprotection)**

The Boc protected side chain (1 eq) was dissolved in dry methanol under a nitrogen atmosphere. 10% Pd/C (0.3 eq) was added followed by ammonium formate (3 eq) and the mixture heated at 75 °C for 30 min. Subsequently, the mixture was cooled to room temperature and the catalyst removed by filtration through a celite pad. The filter cake was washed with methanol and chloroform and the wash fractions were combined with the filtrate. The solvent was removed under reduced pressure and the residue dissolved in dichloromethane and washed with 1 N HCl (aq). The organic layer was dried over $MgSO_4$ and concentrated under reduced pressure.

**Procedure H (side chain coupling to 2)**

The side chain acid (1 eq), HOBt (1 eq), and 1-ethyl-3-(3-dimethylaminopropyl)carbodiimide (EDC, 1.5 eq) were dissolved in DMF (5 mL) and the reaction mixture was stirred at room temperature for 30 min. Subsequently, **2** (1 eq) and *N*,*N*-diisopropylethylamine (4 eq) were added and the mixture stirred over night at room temperature. The mixture was diluted with dichloromethane (8 mL), washed with water, 5% (w/v) LiCl (aq), and brine. The organic layer was dried over $MgSO_4$, concentrated under reduced pressure and the residue purified by column chromatography.

**Procedure I (Boc deprotection)**

The Boc protected intermediate was dissolved in dichloromethane (5 mL), TFA (5 mL) was added and the resultant solution was stirred for 3 h at room temperature. The solvent was removed under reduced pressure, neutralised with saturated $NaHCO_3$ (aq) and extracted with dichloromethane. The organic layer was dried over $MgSO_4$ and concentrated under reduced pressure. The final compounds were either used without further purification or purified by column chromatography.

**Procedure I (Boc deprotection variant b)**

The Boc protected intermediate was dissolved in dichloromethane (5 mL), TFA (5 mL) was added and the resultant solution was stirred for 3 h at room temperature. The deprotected amine was purified using a strong cation exchange (SCX) column. After addition of the reaction mixture, the column was washed with methanol (4 column volumes (CV)), followed by elution with 7 N ammonia in methanol (3 CV). The eluted fractions were concentrated under reduced pressure to yield the final compound.

**Procedure J (acid chloride coupling to 2)**

To a mixture of **2** (1 eq) and triethylamine (6 eq) in dry dichloromethane was added acetyl chloride (1.5 eq). The reaction was stirred at room temperature for 2 h. Subsequently, the solvent was removed under reduced pressure and the residue purified by column chromatography.

**Procedure K (tertiary amine analogue synthesis)**

To a stirred solution of **2** (100 mg, 0.43 mmols) in THF (4 mL) at 0 °C was added DIPEA (150 µL, 0.86 mmol). Chloroacetyl chloride (350 µL, 0.43 mmol) was added dropwise and the reaction allowed to warm to room temperature and stirred for 1 h. The reaction mixture was diluted in dichloromethane and washed with water and brine. The organic layer was dried over $MgSO_4$, filtered and the solvent was removed under reduced pressure to afford a brown oil. The crude material (1 eq crude yield) was taken forward without further purification. A suspension of potassium carbonate (3 eq) and the required amine (1.1 eq) in acetonitrile (2 mL) was added. The reaction mixture was stirred at 60 °C for 16 h. The solvent was removed under reduced pressure and the residue dissolved in EtOAc then washed with water. The organic layer was dried over $MgSO_4$, filtered and the solvent removed under reduced pressure. The residue was dissolved in methanol and purified using an SCX column, eluting with 2 N ammonia in methanol. The solvent was removed under reduced pressure and the residue purified by column chromatography.

**Procedure L (urea synthesis)**

The primary amine (1 eq) was added to a solution of 1,1'-carbonyldiimidazole (1.5 eq) in dichloromethane (1 mL) and the mixture stirred at room temperature for 1 h. **2** (1.3 eq) was added and the reaction mixture was stirred at room temperature overnight. The solvent was removed under reduced pressure and the residue purified by column chromatography.

**Synthesised Compounds**

It should be noted that acylated derivatives of 4,5,6,7-tetrahydrothieno[3,2-*c*]pyridines display two amide rotameric confirmations in NMR spectra.[1]

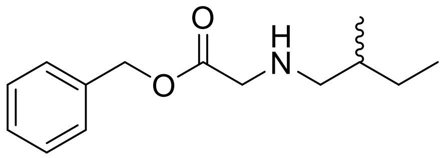

**Benzyl [(2-methylbutyl)amino]acetate (1a).** Compound **1a** was prepared from benzyl bromoacetate (2.8 mmol, 0.44 mL) and 2-methylbutylamine (8.4 mmol, 1.0 mL) using **procedure E** to afford the title compound as a light orange oil (2.2 mmol, 520 mg, 79%). $R_f$ = 0.50 (n-hexane/ EtOAc/NEt$_3$ 50:50:1). $^1$H-NMR (400 MHz, CDCl$_3$) δ (ppm): 7.37-7.34 (m, 5H), 5.17 (s, 2H), 3.45 (s, 2H), 2.51 (dd, *J* = 11 Hz, *J* = 6.0 Hz, 1H), 2.38 (dd, *J* = 11 Hz, *J* = 7.2 Hz, 1H), 1.53-1.36 (m, 2H), 1.20-1.09 (m, 1H), 0.91-0.83 (m, 6H). $^{13}$C-NMR (101 MHz, CDCl$_3$) δ (ppm): 172.6, 135.7, 128.6, 128.4, 66.5, 55.7, 51.3, 35.0, 27.4, 17.6,11.3. HRMS (ESI): calculated for $C_{14}H_{22}NO_2$ [M+H]$^+$: 236.1651, found: 236.1662.

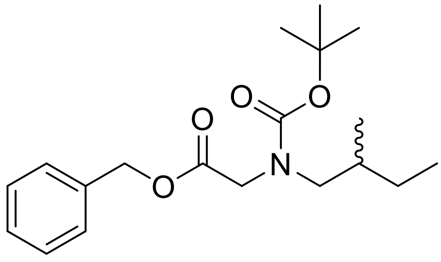

**Benzyl [(*N-tert*-butoxycarbonyl)(2-methylbutyl)amino]acetate (1b).** Compound **1b** was prepared from **1a** (1.5 mmol, 330 mg), triethylamine (2.2 mmol, 0.31 mL), and di-*tert*-butyl dicarbonate (1.5 mmol, 320 mg) using **procedure F** to afford the title compound as an orange oil (1.2 mmol, 390 mg, 81%). $R_f$ = 0.44 (n-hexane/EtOAc 1:4). $^1$H-NMR (400 MHz, CDCl$_3$) δ(ppm): 7.37-7.33 (m, 5H), 5.17 (s, 2H), 4.04-3.94 (m, 1H), 3.89 (s, 1H), 3.21-2.99 (m, 2H), 1.61-1.55 (m, 2H), 1.46-1.35 (m, 10H), 1.13-1.03 (m, 1H), 0.89-8.84 (m, 6H). $^{13}$C-NMR (101 MHz, CDCl$_3$) δ (ppm): 170.0, 135.5, 128.6, 80.1, 66.7, 54.4, 50.0, 34.3, 28.4, 17.0, 11.3. HRMS (ESI): calculated for $C_{19}H_{30}NO_4$ [M+H]$^+$: 336.2175, found: 336.2167.

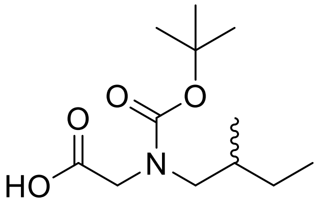

**[(*N-tert*-Butoxycarbonyl)(2-methylbutyl)amino]acetic acid (1c).** Compound **1c** was prepared from **1b** (1.7 mmol, 580 mg), 10% Pd/C (0.10 mmol, 110 mg) and ammonium formate (5.3 mmol, 330 mg) using **procedure G** to afford the title compound as a light brown oil (1.7 mmol, 410 mg, 98%). Characterisation data was consistent with previous reports.[1b]

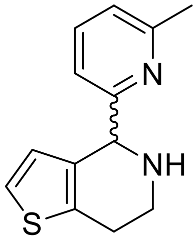

**4-(6-Methylpyridin-2-yl)-4,5,6,7-tetrahydrothieno[3,2-*c*]pyridine (2).** Compound **2** was prepared as reported[1b] or using **Procedure D** from 6-methylpicolinaldehyde (430 mg, 3.5 mmol) to afford the title compound as a brown solid (440 mg, 54% yield). Characterisation data was consistent with previous reports.[1b]

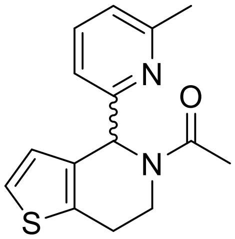

**1-(4-(6-Methylpyridin-2-yl)-6,7-dihydrothieno[3,2-*c*]pyridin-5(4H)-yl)ethan-1-one (3).** Compound **3** was prepared from **2** (0.089 mmol, 20 mg) using **procedure J** to afford the title compound as a yellow oil (0.062 mmol, 17 mg, 95%). $R_f$ = 0.53 (EtOAc) $^1$H-NMR (500 MHz, $CDCl_3$) δ (ppm): 7.56-7.47 (m, 1H), 7.18-7.12 (m, 1H), 7.10-7.04 (m, 1H), 7.03-6.94 (m, 1H), 6.92-6.80 (m, 1H), 6.63 (s, 1H), 6.03 (s, 1H), 5.03-4.95 (m, 1H), 4.09-3.97 (m, 1H), 3.08-2.82 (m, 2H), 2.56 (s, 2H), 2.51 (s, 1H), 2.35 (s, 3H), 2.22 (s, 3H). $^{13}$C-NMR (125 MHz, $CDCl_3$) δ (ppm): 170.8, 159.1, 159.0, 158.6, 135.5, 133.1, 132.8, 126.7, 126.4, 123.2, 122.8, 122.3, 119.0, 117.9, 60.5, 60.4, 55.9, 42.5, 36.6, 29.7, 25.7, 24.9, 24.5, 22.6, 22.2, 21.1, 14.2. HRMS (ESI): calculated for $C_{15}H_{17}N_2O$ $[M+H]^+$: 273.1056, found: 273.1064.

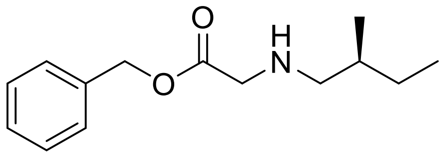

**Benzyl [(*S*)-(2-methylbutyl)amino]acetate (4a).** Compound **4a** was prepared from benzyl bromoacetate (2.8 mmol, 0.44 mL) and (*S*)-2-methylbutylamine (8.4 mmol, 1.0 mL) using **procedure E** to afford the title compound as a colourless oil (2.6 mmol, 620 mg, 93%). $R_f$ = 0.25 (n-hexane/EtOAc/$NEt_3$ 70:30:1). $^1$H-NMR (400 MHz, $CDCl_3$) δ (ppm): 7.37-7.32 (m, 5H), 5.17 (s, 2H), 3.45 (s, 2H), 2.54-2.36 (m, 2H), 1.56-1.37 (m, 12H), 1.20-1.09 (m, 1H), 0.91-0.88 (m, 6H). $^{13}$C-NMR (101 MHz, $CDCl_3$) δ (ppm): 192.9, 172.6, 135.7, 128.6, 128.4, 66.5, 55.7, 51.3, 35.0, 27.4, 17.6, 11.3. HRMS (ESI): calculated for $C_{14}H_{22}NO_2$ $[M+H]^+$: 236.1651, found: 236.1666.

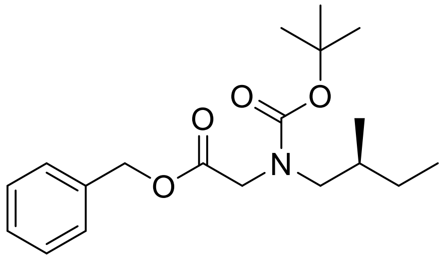

**Benzyl [(*N*-*tert*-butoxycarbonyl)-(*S*)-(2-methylbutyl)amino]acetate (4b).** Compound **4b** was prepared from **4a** (2.2 mmol, 520 mg), triethylamine (3.30 mmol, 0.46 mL) and di-*tert*-butyl dicarbonate (2.2 mmol, 480 mg) using **procedure F** to afford the title compound as a colourless oil (1.9 mmol, 620 mg, 85%). $R_f$ = 0.45 (n-hexane/EtOAc 1:4). $^1$H-NMR (400 MHz, $CDCl_3$) δ(ppm): 7.42-7.31 (m, 5H), 5.19 (d, *J* = 1.7 Hz, 2H), 4.09-3.75 (m, 2H), 3.26-2.99 (m, 2H), 1.67-1.56 (m, 2H), 1.51-1.33 (m, 9H), 1.07-1.05 (m, 1H), 0.96-0.84 (m, 6H). $^{13}$C-NMR (101 MHz, $CDCl_3$) δ (ppm): 128.6, 128.5, 128.4, 128.3,128.2, 66.7, 66.7, 54.4, 54.4, 50.0, 49.4, 34.3, 34.0, 28.4, 28.3, 28.2, 26.9, 17.0, 16.8, 11.3. HRMS (ESI): calculated for $C_{19}H_{30}NO_4$ $[M+H]^+$: 336.2178, found: 336.2178.

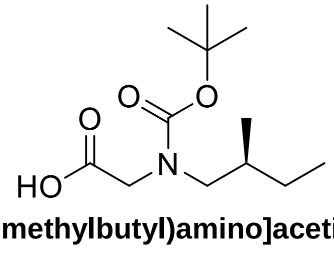

**[(*N*-*tert*-Butoxycarbonyl)-(*S*)-(2-methylbutyl)amino]acetic acid (4c).** Compound **4c** was prepared from **4b** (1.9 mmol, 620 mg), 10% Pd/C (0.23 mmol, 240 mg) and ammonium formate (4.6 mmol, 290 mg) using **procedure G** to afford the title compound as a brown oil (1.9 mmol, 460 mg, 100%). $^1$H-NMR (400 MHz, $CDCl_3$) δ (ppm): 4.04-3.82 (m, 2H), 3.30-2.89 (m, 2H), 1.66-1.52 (m, 1H), 1.52-1.22 (m, 10H), 1.10 (m, 1H), 0.88 (q, *J* = 6.6, 5.8 Hz, 6H). $^{13}$C-NMR (101 MHz, $CDCl_3$) δ (ppm): 54.6, 54.2, 49.5, 34.2, 34.0, 28.3, 28.2, 26.9, 17.0, 16.8, 11.3. HRMS (ESI): calculated for $C_{12}H_{24}NO_4$ $[M+H]^+$: 246.1705, found: 246.1693.

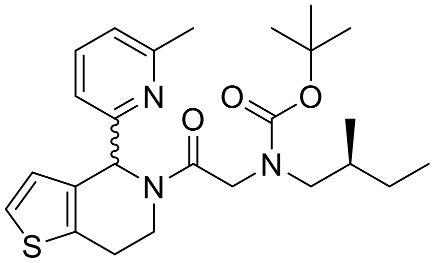

***tert*-Butyl (*S*)-(2-methylbutyl)(2-(4-(6-methylpyridin-2-yl)-6,7-dihydrothieno[3,2-*c*]pyridin-5(4*H*)-yl)-2-oxoethyl)carbamate** (**4d**). The Boc-protected intermediate **4d** was prepared **4c** (0.15 mmol, 37 mg) and **2** (0.18 mmol, 42 mg) using **procedure H** to afford the title compound as a colourless oil (0.12 mmol, 57 mg, 80%). $R_f$ = 0.3 (n-hexane/EtOAc 3:7). $^1$H-NMR (400 MHz, $CDCl_3$) δ (ppm): 7.48 (q, J = 7.6 Hz, 1H), 7.23-7.11 (m, 1H), 7.10-6.95 (m, 1H), 6.92-6.74 (m, 2H), 6.53 (s, 1H), 5.98 (d, J = 15 Hz, 1H), 4.95 (d, J = 11 Hz, 1H), 4.88 (d, J = 11 Hz, 1H), 4.74-4.47 (m, 1H), 4.43-4.21 (m, 1H), 4.16-3.94 (m, 1H), 3.43 (dd, J = 15, 6.6 Hz, 1H), 3.31-2.72 (m, 4H), 2.60-2.41 (m, 3H), 1.53-1.30 (m, 9H), 1.16-0.98 (m, 1H), 0.94-0.75 (m, 6H). HRMS (ESI): calculated for $C_{25}H_{36}N_3O_3S$ $[M+H]^+$: 458.2477, found: 458.2490.

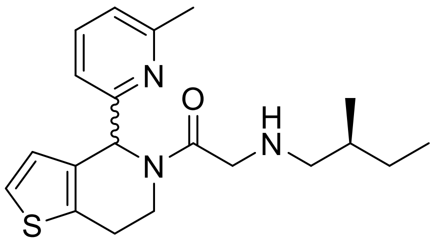

**2-(*S*)-(2-Methylbutylamino)-1-[4-(6-methylpyridin-2-yl)-6,7-dihydro-4*H*-thieno[3,2-*c*]pyridin-5-yl]ethanone** (**4**). Compound **4** was prepared from the Boc-protected intermediate **4d** (0.11 mmol, 52 mg) using **procedure I** to afford the title compound as a yellow oil (0.90 mmol, 32 mg, 79%). $R_f$ = 0.17 (EtOAc). Chiral HPLC (**Method A**): RT = 9.1 min (50%), 12.7 min (50%). $^1$H-NMR (500 MHz, $CDCl_3$) δ (ppm): 7.49 (q, J = 7.4 Hz, 1H), 7.14 (d, J = 5.8 Hz, 1H), 7.09-6.94 (m, 1H), 6.92-6.77 (m, 1H), 6.56 (s, 1H), 5.98 (s, 1H), 4.94 (dd, J = 12, 4.0 Hz, 1H), 4.06-3.97 (m, 1H), 3.71 (dd, J = 16, 5.4 Hz, 1H), 3.56 (qd, J = 16, 5.7 Hz, 1H), 3.09-2.79 (m, 1H), 2.58-2.49 (m, 1H), 2.47-2.39 (m, 2H), 1.61-1.49 (m, 1H), 1.48-1.36 (m, 1H), 1.14 (m, 1H), 0.94-0.82 (m, 6H). $^{13}$C-NMR (125 MHz, $CDCl_3$) δ (ppm): 171.1, 169.9, 159.0, 158.6, 158.4, 136.8, 136.6, 135.5, 134.1, 132.9, 132.3, 126.6, 126.2, 123.3, 122.8, 122.3, 121.9, 118.8, 118.2, 58.8, 56.4, 56.3, 56.1, 51.4, 51.2, 40.6, 36.8, 35.0, 35.0, 34.8, 27.4, 27.4, 27.3, 25.6, 24.8, 24.6, 24.4, 17.6, 17.5, 11.3, 11.2, 11.2. HRMS (ESI): calculated for $C_{20}H_{28}N_3OS$ $[M+H]^+$: 358.1953, found: 358.1960.

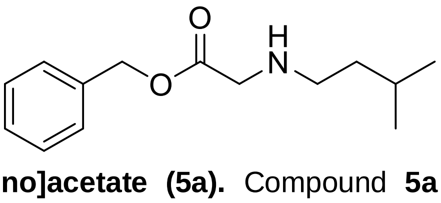

**Benzyl [(3-methylbutyl)amino]acetate (5a).** Compound **5a** was prepared from benzyl bromoacetate (2.8 mmol, 0.44 mL) and isopentylamine (8.4 mmol, 1.0 mL) using **procedure**

**E** to afford the title compound as a colourless oil (0.73 mmol, 170 mg, 25%). $R_f$ = 0.27 (n-hexane/ EtOAc/NEt$_3$ 70:30:1). $^1$H-NMR (400 MHz, CDCl$_3$) δ (ppm): 7.41-7.29 (m, 5H), 5.17 (s, 2H), 3.46 (s, 2H), 2.65-2.54 (m, 2H), 1.63 (m, 1H), 1.43-1.31 (m, 2H), 0.89 (d, *J* = 6.8 Hz, 6H). $^{13}$C-NMR (101 MHz, CDCl$_3$) δ (ppm): 172.5, 135.6, 128.6, 128.4, 66.5, 51.0, 47.7, 39.1, 26.0, 22.3. HRMS (ESI): calculated for: $C_{14}H_{22}NO_2$ [M+H]$^+$: 236.1651, found: 236.1653.

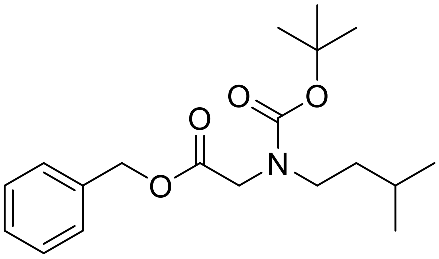

**Benzyl [(*N*-*tert*-butoxycarbonyl)(3-methylbutyl)amino]acetate (5b).** Compound **5b** was prepared from **5a** (0.73 mmol, 170 mg), triethylamine (1.1 mmol, 0.16 mL) and di-*tert*-butyl dicarbonate (0.73 mmol, 160 mg) using **procedure F** to afford the title compound as a colourless oil (0.53 mmol, 180 mg, 73%). $R_f$ = 0.40 (n-hexane/EtOAc 1:4). $^1$H-NMR (400 MHz, CDCl$_3$) δ(ppm): 7.35 (s, 5H), 5.17 (s, 2H), 3.99 (s, 1H), 3.88 (s, 1H), 3.27 (dt, *J* = 21, 7.7 Hz, 2H), 1.61-1.33 (m, 11H), 0.89 (d, *J* = 6.5 Hz, 7H). $^{13}$C-NMR (101 MHz, CDCl$_3$) δ (ppm): 170.0, 128.6, 128.5, 128.4, 128.3, 80.0, 66.7, 49.3, 48.7, 46.8, 37.3, 36.9, 28.4, 28.2, 27.4, 25.9, 25.6, 22.5. HRMS (ESI): calculated for $C_{19}H_{30}NO_4$ [M+H]$^+$: 336.2175, found: 336.2180.

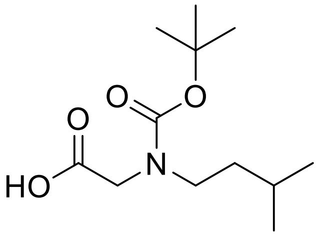

**[(*N*-*tert*-Butoxycarbonyl)(3-methylbutyl)amino]acetic acid (5c).** Compound **5c** was prepared from **5b** (0.52 mmol, 180 mg), 10% Pd/C (0.16 mmol, 100 mg) and ammonium formate (3.1 mmol, 190 mg) using **procedure L** to afford the title compound as a brown oil (0.46 mmol, 110 mg, 88%). $^1$H-NMR (400 MHz, CDCl$_3$) δ (ppm): 3.97 (s, 1H), 3.90 (s, 1H), 3.28 (m, 2H), 1.63-1.29 (m, 12H), 0.91 (d, *J* = 6.6 Hz, 6H). $^{13}$C-NMR (101 MHz, CDCl$_3$) δ (ppm): 77.3, 77.0, 76.7, 49.1, 47.1, 37.2, 28.3, 25.6, 22.5. HRMS (ESI): calculated for $C_{12}H_{22}NO_4$ [M+H]$^+$: 244.1549, found: 244.1553.

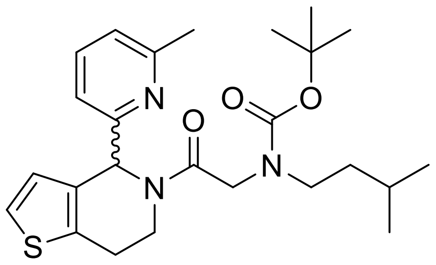

***tert*-Butyl isopentyl(2-(4-(6-methylpyridin-2-yl)-6,7-dihydrothieno[3,2-*c*]pyridin-5(4*H*)-yl)-2-oxoethyl)carbamate (5d).** The Boc-protected intermediate **5d** was prepared **5c** (0.096 mmol, 25 mg) and **2** (0.096 mmol, 22 mg) using **procedure H** to afford the title compound as a colourless oil (0.066 mmol, 28 mg, 60%). $R_f$ = 0.3 (n-hexane/EtOAc 3:7). $^1$H-NMR (400 MHz, $CDCl_3$) δ (ppm): 7.49 (q, *J* = 8.0 Hz, 2H), 7.22-7.10 (m, 2H), 7.05 (t, *J* = 7.1 Hz, 2H), 6.98 (d, *J* = 7.7 Hz, 1H), 6.65-6.94 (m, 3H), 6.53 (s, 1H), 6.02 (s, 1H), 5.97 (s, 1H), 4.99-4.83 (m, 1H), 4.58 (d, *J* = 17 Hz, 1H), 4.38-4.27 (m, 1H), 4.13-3.98 (m, 2H), 3.44 (dt, *J* = 15 Hz, *J* = 7.8 Hz, 1H), 3.10-3.35 (m, 2H), 3.12-2.76 (m, 5H), 2.55 (s, 3H), 2.45 (s, 2H), 1.59-1.50 (m, 1H), 1.49-1.23 (m, 14H), 1.23-1.11 (m, 1H), 0.93-0.83 (m, 10H). $^{13}$C-NMR (101 MHz, $CDCl_3$) δ (ppm): 168.7, 158.9, 158.6, 136.8, 126.6, 126.2, 123.3, 122.3, 121.9, 118.9, 118.4, 79.5, 59.0, 58.7, 56.5, 49.4, 48.7, 46.6, 46.3, 41.1, 37.1, 36.9, 36.5, 29.7, 28.4, 26.1, 25.8, 24.9, 24.5, 22.6, 22.6. HRMS (ESI): calculated for $C_{25}H_{36}N_3O_3S$ $[M+H]^+$: 458.2472, found: 458.2462.

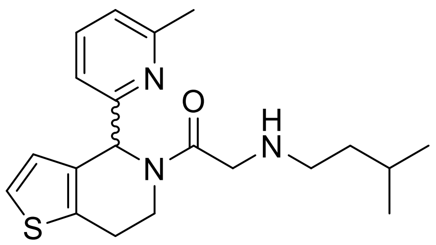

**2-(Isopentylamino)-1-(4-(6-methylpyridin-2-yl)-6,7-dihydrothieno[3,2-*c*]pyridin-5(4*H*)-yl)ethan-1-one (5).** Compound **5** was prepared from the Boc-protected intermediate **5d** (0.045 mmol, 19 mg) using **procedure I** to afford the title compound as a colourless oil (0.045 mmol, 15 mg, 100%). $R_f$ = 0.18 (n-hexane/EtOAc 3:1). $^1$H-NMR (500 MHz, $CDCl_3$) δ (ppm): 7.53-7.45 (m, 1H), 7.14 (d, *J* = 5.2 Hz, 1H), 7.10-7.02 (m, 1H), 6.98 (d, *J* = 7.7 Hz, 1H), 6.90 (d, *J* = 5.2 Hz, 1H), 6.85 (d, *J* = 5.2 Hz, 1H), 6.80 (d, *J* = 7.7 Hz, 1H), 6.56 (d, *J* = 1.5 Hz, 1H), 5.97 (s, 1H), 4.98-4.90 (m, 1H), 4.09-3.94 (m, 2H), 3.70 (d, *J* = 16 Hz, 1H), 3.57 (q, *J* = 16 Hz, 1H), 3.08-2.83 (m, 3H), 2.69-2.57 (m, 3H), 2.54 (s, 2H), 2.45 (s, 1H), 1.60-1.65 (m, 1H), 1.46-1.36 (m, 2H), 1.23-1.12 (m, 1H), 0.92-0.85 (m, 6H). $^{13}$C-NMR (125 MHz, $CDCl_3$) δ (ppm): 171.0, 169.8, 159.0, 158.6, 158.4, 136.8, 136.6, 135.6, 134.1, 132.9, 132.2, 126.6, 126.2, 123.3, 122.8, 122.4, 121.8, 118.8, 118.2, 58.8, 56.4, 51.1, 50.9, 50.6, 48.3, 48.1, 40.6, 39.0, 38.8, 36.8, 26.0, 25.6, 24.8, 24.6, 24.5, 22.6, 22.6. HRMS (ESI): calculated for $C_{20}H_{28}N_3OS$ $[M+H]^+$: 358.1948, found: 358.1968.

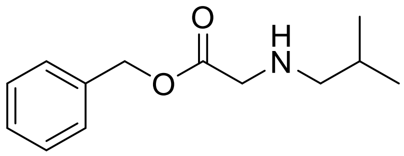

**Benzyl [(2-methylpropyl)amino]acetate (6a).** Compound **6a** was prepared from benzyl bromoacetate (2.4 mmol, 0.53 mL) and isobutylamine (11 mmol, 1.0 mL) using **procedure E** to afford the title compound as a colourless oil (2.4 mmol, 530 mg, 71%). $R_f$ = 0.29 (n-hexane/ EtOAc/$NEt_3$ 70:30:1). $^1$H-NMR (400 MHz, $CDCl_3$) δ (ppm): 7.42-7.28 (m, 5H), 5.17 (s, 2H), 3.45 (d, *J* = 0.8 Hz, 2H), 2.42 (d, *J* = 6.8 Hz, 2H), 1.73 (m, 1H), 0.91 (d, *J* = 6.8 Hz, 6H). $^{13}$C-NMR (101 MHz, $CDCl_3$) δ (ppm): 172.5, 135.7, 128.6, 128.4, 66.5, 57.6, 51.2, 28.5, 20.6. HRMS (ESI): calculated for $C_{13}H_{20}NO_2$ $[M+H]^+$: 222.1494, found: 222.1503.

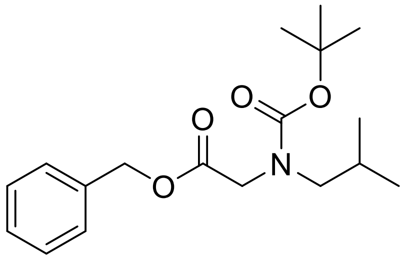

**Benzyl [(*N-tert*-butoxycarbonyl)(2-methylpropyl)amino]acetate (6b).** Compound **6b** was prepared from **6a** (2.3 mmol, 500 mg), triethylamine (3.5 mmol, 0.48 mL) and di-*tert*-butyl dicarbonate (2.3 mmol, 500 mg) using **procedure F** to afford the title compound as a colorless oil (2.1 mmol, 670 mg, 90%). $R_f$ = 0.28 (n-hexane/EtOAc 1:4). $^1$H-NMR (400 MHz, $CDCl_3$) δ(ppm): 7.42-7.31 (m, 5H), 5.22 – 5.16 (m, 2H), 4.02 (s, 1H), 3.92 (s, 1H), 3.11 (dd, *J* = 18, 7.4 Hz, 2H), 1.84 (m, 1H), 1.43-1.38 (m, 9H), 0.90 (t, *J* = 6.5 Hz, 6H). $^{13}$C-NMR (101 MHz, $CDCl_3$) δ (ppm): 128.6, 128.5, 128.4, 128.3, 128.2, 80.0, 66.7, 66.7, 56.0, 55.8, 50.1, 49.4, 28.3, 28.2, 27.9, 27.6, 20.1, 20.0, 14.2. HRMS (ESI): calculated for $C_{18}H_{28}NO_4$ $[M+H]^+$: 322.2018, found: 322.2026.

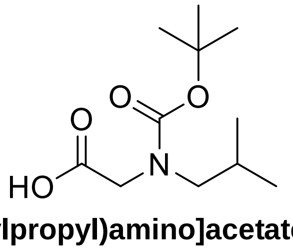

**[(*N-tert*-Butoxycarbonyl)(2-methylpropyl)amino]acetate (6c).** Compound **6c** was obtained from **6b** (2.8 mmol, 640 mg), 10% Pd/C (0.8 mmol, 880 mg) and ammonium formate (16 mmol, 1.0 g) using **procedure G** to afford the title compound as a brown oil (1.2 mmol, 370 mg, 42%). $^1$H-NMR (400 MHz, $CDCl_3$) δ (ppm): 10.75 (s, 1H), 4.00 (s, 1H), 3.91 (s, 1H), 3.10 (dd, *J* = 13, 7.3 Hz, 2H), 1.84 (m, 1H), 1.46 (d, *J* = 16 Hz, 9H), 0.90 (d, *J* = 6.7 Hz, 6H). $^{13}$C-NMR (101 MHz, $CDCl_3$) δ (ppm): 80.9, 80.4, 56.2, 55.7, 49.6, 29.7, 28.3, 28.2, 27.8, 27.6, 20.0. HRMS (ESI): calculated for $C_{11}H_{20}NO_4$ $[M+H]^+$: 230.1392, found: 230.1399.

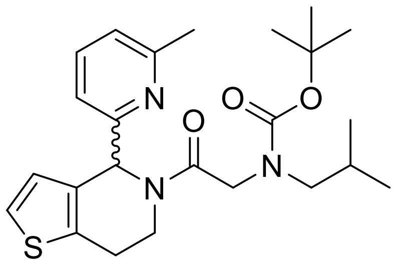

***tert*-Butyl isobutyl(2-(4-(6-methylpyridin-2-yl)-6,7-dihydrothieno[3,2-*c*]pyridin-5(4*H*)-yl)-2-oxoethyl)carbamate (6d).** The Boc-protected intermediate **6d** was prepared from **6c** (0.090 mmol, 20 mg) and **2** (0.090 mmol, 20 mg) using **procedure H** to afford the title compound as a colourless oil (0.080 mmol, 37 mg, 89%). $R_f$ = 0.23 (n-hexane/EtOAc 3:7). $^1$H-NMR (400 MHz, $CDCl_3$) δ (ppm): 7.49 (q, *J* = 7.7 Hz, 1H), 7.22-6.94 (m, 2H), 6.75-6.92 (m, 2H), 6.53 (s, 1H), 5.98 (d, *J* = 17 Hz, 1H), 4.98-4.83 (m, 1H), 4.62 (q, *J* = 17 Hz, 1H), 4.47-4.26 (m, 1H), 4.17-3.89 (m, 1H), 3.34-2.76 (m, 5H), 2.57-2.39 (m, 3H), 1.94-1.79 (m, 2H), 1.51-1.25 (m, 9H), 0.94-0.81 (m, 6H). $^{13}$C-NMR (101 MHz, $CDCl_3$) δ (ppm): 158.9, 158.6, 156.5, 136.8, 135.9, 126.1, 123.3, 122.4, 118.6, 79.5, 59.0, 58.6, 56.5, 55.6, 50.1, 49.4, 36.8, 36.4, 29.7, 28.4, 28.3, 27.6, 27.4, 25.7, 24.9, 24.5, 20.1, 14.2. HRMS (ESI): calculated for: $C_{24}H_{34}N_3O_3S$ $[M+H]^+$: 444.2315, found: 444.2323.

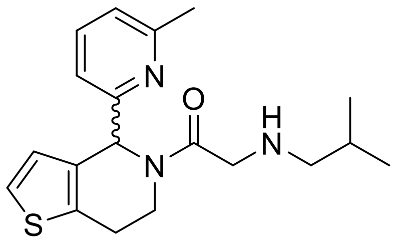

**2-(Isobutylamino)-1-(4-(6-methylpyridin-2-yl)-6,7-dihydrothieno[3,2-*c*]pyridin-5(4*H*)-yl)ethan-1-one (6).** Compound **6** was prepared from Boc-protected intermediate **6d** (0.080 mmol, 37 mg) using **procedure I** to afford the title compound as a colourless oil (0.080 mmol, 28 mg, 100%). Chiral HPLC (Method A): room temperature = 9.6 min (50%), 13.8 min (50%). $^1$H-NMR (500 MHz, $CDCl_3$) δ (ppm): 7.50 (q, *J* = 7.6 Hz, 1H), 7.15 (t, J = 5.7 Hz, 1H), 7.06 (dd, *J* = 12 Hz, *J* = 6.4 Hz, 1H), 6.98 (d, *J* = 7.7 Hz, 1H), 6.89 (d, *J* = 5.2 Hz, 1H), 6.85 (d, *J* = 5.2 Hz, 1H), 6.80 (d, *J* = 7.7 Hz, 1H), 6.56 (d, *J* = 1.5 Hz, 1H), 5.97 (s, 1H), 4.98-4.87 (m, 1H), 4.12 (d, *J* = 16 Hz, 1H), 4.06-3.96 (m, 1H), 3.76 (d, *J* = 16 Hz, 1H), 3.59 (q, *J* = 16 Hz, 1H), 3.07-2.81 (m, 1H), 2.54 (s, 2H), 2.50-2.39 (m, 3H), 1.86-1.73 (m, 1H), 0.98-0.90 (m, 6H). $^{13}$C-NMR (125 MHz, $CDCl_3$) δ (ppm): 170.6, 169.7, 158.9, 158.7, 158.5, 158.4, 136.9, 136.6, 135.6, 134.1, 132.9, 132.1, 126.6, 126.2, 123.4, 122.8, 122.4, 121.9, 118.8, 118.3, 58.7, 58.1, 57.9, 56.5, 51.2, 50.9, 40.7, 36.8, 28.5, 28.2, 25.6, 24.8, 24.6, 24.5, 20.6, 20.6, 20.5. HRMS (ESI): calculated for $C_{19}H_{26}N_3OS$ $[M+H]^+$: 344.1791, found: 344.1796.

**(-)-2-(Isobutylamino)-1-(4-(6-methylpyridin-2-yl)-6,7-dihydrothieno[3,2-*c*]pyridin-5(4*H*)-yl)ethan-1-one ((-)-6).** Compound **(-)-6** was obtained from compound **6** by preparative chiral HPLC (**Method D**) (*er* 100:0). Chiral HPLC (**Method A**): 13.5 min (100%). $[\alpha]^{23}_D$ (c = 0.1, $CHCl_3$): -57.2. $^1$H-NMR (500 MHz, $CDCl_3$) δ (ppm): 7.50 (q, *J* = 7.6 Hz, 1H), 7.32 (d, *J* = 5.7 Hz, 1H), 7.10-7.02 (m, 1H), 6.98 (d, *J* = 7.7 Hz, 1H), 6.90 (d, *J* = 5.2 Hz, 1H), 6.85 (d, *J* = 5.2 Hz,

1H), 6.81 (d, *J* = 7.7 Hz, 1H), 6.56 (s, 1H), 5.98 (s, 1H), 4.99-4.91 (m, 1H), 4.08-3.96 (m, 2H), 3.70 (d, *J* = 16 Hz, 1H), 3.56 (q, *J* = 16 Hz, 1H), 3.09-2.83 (m, 3H), 2.54 (s, 2H), 2.50-2.37 (m, 3H), 1.82-1.71 (m, 1H), 0.95-0.81 (m, 6H). $^{13}$C-NMR (125 MHz, $CDCl_3$) δ (ppm): 171.1, 170.0, 159.0, 158.7, 158.4, 136.8, 136.6, 135.6, 134.1, 132.9, 132.3, 126.6, 126.3, 123.4, 122.8, 122.4, 121.9, 118.8, 118.2, 58.8, 58.2, 58.0, 56.4, 51.4, 51.1, 40.6, 36.8, 29.7, 28.6, 28.4, 25.6, 24.8, 24.6, 24.5, 20.7, 20.6, 20.6. HRMS (ESI): calculated for $C_{19}H_{26}N_3OS$ $[M+H]^+$: 344.1791, found: 344.1800.

**(+)-2-(Isobutylamino)-1-(4-(6-methylpyridin-2-yl)-6,7-dihydrothieno[3,2-*c*]pyridin-5(4*H*)-yl)ethan-1-one ((+)-6, IMP-1575).** Compound **(+)-6** was obtained from compound **6** by preparative chiral HPLC (**Method D**) (*er* 100:0). Chiral HPLC (**Method A**): 9.4 min (100%). $[\alpha]^{23}_D$ (c = 0.67, $CHCl_3$): +43.6. $^1$H-NMR (500 MHz, $CDCl_3$) δ (ppm): 7.50 (q, *J* = 7.6 Hz, 1H), 7.17-7.12 (m, 1H), 7.10-7.02 (m, 1H), 6.98 (d, *J* = 7.7 Hz, 1H), 6.90 (d, *J* = 5.2 Hz, 1H), 6.85 (d, *J* = 5.2 Hz, 1H), 6.80 (d, *J* = 7.7 Hz, 1H), 6.56 (d, *J* = 1.5 Hz, 1H), 5.97 (s, 1H), 5.00-4.90 (m, 1H), 4.11-3.97 (m, 1.7H), 3.73 (d, *J* = 16.2 Hz, 1H), 3.57 (q, *J* = 16 Hz, 1H), 3.08-2.83 (m, 3H), 2.54 (s, 2H), 2.52-2.38 (m, 3H), 1.84-1.71 (m, 1H), 0.96-0.84 (m, 6H). $^{13}$C-NMR (125 MHz, $CDCl_3$) δ (ppm): 170.9, 169.9, 159.0, 158.7, 158.6, 158.4, 136.9, 136.6, 135.6, 134.1, 132.9, 132.2, 126.6, 126.2, 123.4, 122.8, 122.4, 121.9, 118.8, 118.2, 58.8, 58.2, 57.9, 56.5, 51.3, 51.0, 40.6, 36.8, 29.7, 28.6, 28.3, 25.6, 24.9, 24.6, 24.5, 20.7, 20.6, 20.6. HRMS (ESI): calculated for $C_{19}H_{26}N_3OS$ $[M+H]^+$: 344.1791, found: 344.1811.

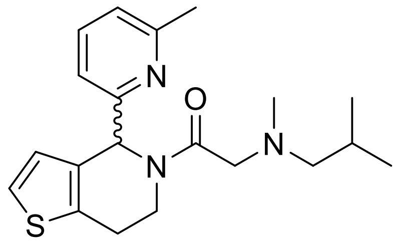

**2-(Isobutyl(methyl)amino)-1-(4-(6-methylpyridin-2-yl)-6,7-dihydrothieno[3,2-*c*]pyridin-5(4*H*)-yl)ethan-1-one (7).** The required tertiary amine was prepared from **2** via the chloroacetamide (66 mg crude yield, 0.22 mmol) and *N*-methyl isobutylamine (29 µL, 0.24 mmol) using **procedure K** to afford the title compound as a yellow oil (0.022 mmol, 7.9 mg, 10%). $R_f$ = 0.25 (n-hexane/EtOAc 1:1.5 with 1% triethylamine) $^1$H-NMR (400 MHz, $CDCl_3$) δ (ppm): 7.52-7.41 (m, 1H), 7.16-7.08 (m, 1H), 7.06 (d, *J* = 5.3 Hz, 1H), 7.01 (d, *J* = 7.7 Hz, 1H), 6.97 (d, *J* = 7.7 Hz, 1H), 6.89 (t, *J* = 5.0 Hz, 1H), 6.85 (s, 1H), 6.79 (d, *J* = 7.7 Hz, 1H), 6.55 (d, *J* = 1.6 Hz, 1H), 4.96-4.85 (m, 1H), 4.54 (ddd, *J* = 13.2, 5.4, 1.7 Hz, 1H), 3.96 (d, *J* = 13.2 Hz, 1H), 3.90 (ddd, *J* = 13.3, 11.8, 3.9 Hz, 1H), 3.33-3.17 (m, 1H), 3.15 (d, *J* = 13.3 Hz, 1H), 3.10-2.76 (m, 3H), 2.52 (s, 2H), 2.45 (s, 1H), 2.23 (s, 2H), 2.20 (s, 1H), 2.17-2.13 (m, 1H), 2.11 (dd, *J* = 7.4, 1.4 Hz, 1H), 1.79-1.63 (m, 1H), 0.85 (t, *J* = 6.9 Hz, 3H), 0.82 (d, *J* = 6.6 Hz, 2H), 0.72 (d, *J* = 6.5 Hz, 2H). $^{13}$C-NMR (101 MHz, $CDCl_3$) δ (ppm): 170.4, 169.3, 159.3, 159.1, 158.3, 158.1, 136.4, 136.3, 134.8, 134.0, 133.3, 133.1, 126.6, 126.3, 122.7, 122.3, 121.7, 121.5, 118.5, 117.9, 66.2, 65.9, 63.2, 62.6, 58.4, 56.0, 42.4, 41.83, 41.4, 36.2, 25.8, 24.8, 24.4, 24.3, 20.7, 20.6, 20.4, 20.2. HRMS (ESI): calculated for $C_{20}H_{28}N_3OS$ $[M+H]^+$: 358.1953, found: 358.1962.

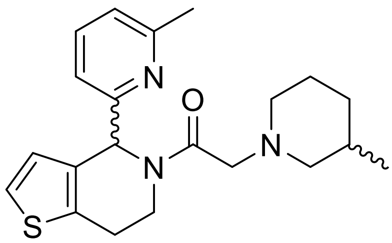

**2-(3-Methylpiperidin-1-yl)-1-(4-(6-methylpyridin-2-yl)-6,7-dihydrothieno[3,2-*c*]pyridin-5(4*H*)-yl)ethan-1-one (8).** The required tertiary amine was prepared from **2** via the chloroacetamide (66 mg crude yield, 0.22 mmol) and 3-methyl piperidine (28 µL, 0.24 mmol) using **Procedure K** to afford the title compound as a yellow oil (0.024 mmol, 9.0 mg, 11%). $R_f$ = 0.32 (n-hexane/EtOAc 1:1.5 with 1% triethylamine). $^1$H NMR (400 MHz, $CDCl_3$) δ (ppm): 7.51-7.43 (m, 2H), 7.12 (d, *J* = 5.5 Hz, 2H), 7.06 (dd, *J* = 5.4, 2.1 Hz, 1H), 7.03-6.94 (m, 4H), 6.92 (dd, *J* = 5.2, 1.1 Hz, 1H), 6.83 (dd, *J* = 7.7, 2.2 Hz, 1H), 6.58 (s, 1H), 6.54-6.50 (m, 2H), 4.92 (dt, *J* = 13, 3.4 Hz, 1H), 4.49-4.37 (m, 1H), 3.94-3.80 (m, 1H), 3.65 (dd, *J* = 14, 11 Hz, 1H), 3.32 (dd, *J* = 16, 14 Hz, 2H), 3.18 (d, *J* = 4.2 Hz, 1H), 3.15 (d, *J* = 4.2 Hz, 1H), 3.10-2.96 (m, 1H), 2.96-2.86 (m, 2H), 2.86-2.67 (m, 5H), 2.62-2.59 (m, 1H), 2.52 (s, 3H), 2.46 (s, 1H), 2.45 (s, 2H), 2.15 (d, *J* = 7.5 Hz, 1H), 1.98 (d, *J* = 18 Hz, 3H), 1.88 (s, 1H), 1.78-1.20 (m, 6H),0.87-0.79 (m, 4H), 0.77 (dd, *J* = 6.7, 2.2 Hz, 3H). $^{13}$C NMR (101 MHz, $CDCl_3$) δ (ppm): 170.2, 169.4, 159.4, 158.6, 158.4, 136.6, 135.2, 134.3, 133.4, 127.0, 126.7, 123.0, 122.6, 121.98, 121.8, 118.7, 118.0, 63.1, 62.4, 62.3, 62.1, 62.0, 61.7, 59.2, 56.4, 54.3, 54.2, 53.8, 42.0, 41.3, 37.00, 32.6, 31.9, 31.4, 31.2, 30.9, 30.6, 26.14, 25.7, 25.49, 25.4, 25.2, 25.1, 24.7, 19.7, 19.5. HRMS (ESI): calculated for $C_{21}H_{28}N_3OS$ $[M+H]^+$: 370.1953, found: 370.1949.

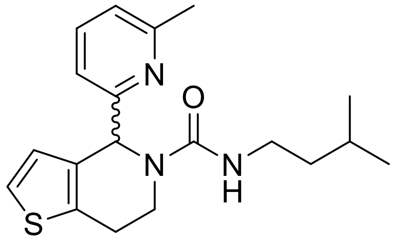

***N*-Isopentyl-4-(6-methylpyridin-2-yl)-6,7-dihydrothieno[3,2-*c*]pyridine-5(4*H*)-carboxamide (9).** Compound **9** was prepared from isopentylamine (0.076 mmol, 7 mg) and **2** (0.096 mmol, 22 mg) using **procedure L** to afford the title compound as a colourless oil (0.048 mmol, 13 mg, 64%). $R_f$ = 0.2 (n-hexane/EtOAc 3:7). $^1$H-NMR (400 MHz, $CDCl_3$) δ (ppm): 7.52 (t, *J* = 7.7 Hz, 1H), 7.21 (t, *J* = 5.2 Hz, 1H), 7.13-7.06 (m, 2H), 6.77 (d, *J* = 7.7 Hz, 1H), 6.61 (d, *J* = 5.2 Hz, 1H), 5.88 (s, 1H), 4.64-4.55 (m, 1H), 3.33-3.20 (m, 2H), 3.06-2.79 (m, 3H), 2.56 (s, 3H), 1.71 (q, *J* = 6.7 Hz, 1H), 1.56-1.39 (m, 2H), 0.94 (d, *J* = 6.7 Hz, 6H). $^{13}$C-NMR (101 MHz, $CDCl_3$) δ (ppm): 159.9, 158.9, 157.6, 137.4, 136.7, 132.1, 125.8, 123.0, 122.6, 119.0, 59.4, 39.1, 39.1, 37.1, 25.8, 25.3, 24.2, 22.6, 22.6. HRMS (ESI): calculated for $C_{19}H_{26}N_3OS$ $[M+H]^+$: 344.1791, found: 344.1781.

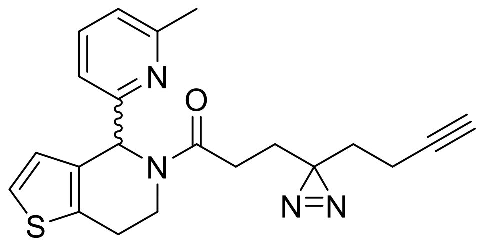

**3-(3-(But-3-yn-1-yl)-3*H*-diazirin-3-yl)-1-(4-(6-methylpyridin-2-yl)-6,7-dihydrothieno[3,2-*c*]pyridin-5(4*H*)-yl)propan-1-one (10).** 3-(3-(But-3-yn-1-yl)-3H-diazirin-3-yl)propanoic acid was synthesised according to known literature procedures.[2] Probe **10** was prepared from **2** (18 mg, 0.078 mmol, 1.0 eq.) and 3-(3-(but-3-yn-1-yl)-3H-diazirin-3-yl)propanoic acid (13 mg, 0.078 mmol, 1.0 eq.) using **procedure H** to afford a crude product that was purified by preparative HPLC-MS to afford the title compound (0.021 mmol, 8 mg, 27 %) $R_f = 0.87$ (EtOAc/n-hexane 20:1). $^1$H NMR (500 MHz, $CDCl_3$) δ(ppm): 7.50 (t, *J* = 8.0 Hz, 1H), 7.49 (t, *J* = 8.0 Hz, 1H), 7.14 (d, *J* = 5.2 Hz, 1H), 7.13 (d, *J* = 8.0 Hz, 1H), 7.08 (d, *J* = 5.2 Hz, 1H), 7.06 (d, *J* = 8.0 Hz, 1H), 6.97 (d, *J* = 7.5 Hz, 1H), 6.91 (d, *J* = 5.2 Hz, 1H), 6.85 (d, *J* = 5.2 Hz, 1H), 6.80 (d, *J* = 7.5 Hz, 1H), 6.55 (s, 1H), 6.01 (s, 1H), 4.97-4.90 (m, 1H), 4.06-3.99 (m, 1H), 3.02-2.90 (m, 2H), 2.89-2.81 (m, 1H), 2.76 (dt, *J* = 16, 7.7 Hz, 1H), 2.57 (s, 3H), 2.45 (s, 3H), 2.36 (dt, *J* = 16, 7.7 Hz, 1H), 2.28-2.11 (m, 2H), 2.02 (td, *J* = 7.6, 2.6 Hz, 2H), 1.96 (t, *J* = 2.6 Hz, 1H), 1.94 (t, *J* = 2.6 Hz, 1H), 1.89 (t, *J* = 7.7 Hz, 2H), 1.84 (t, *J* = 7.7 Hz,, 2H), 1.66 (t, *J* = 7.5 Hz, 2H), 1.66 (t, *J* = 7.5 Hz, 2H). $^{13}$C-NMR (126 MHz, $CDCl_3$) δ (ppm): 171.5, 170.0, 159.2, 159.0, 158.8, 158.5, 137.0, 136.8, 135.8, 134.4, 132.8, 132.7, 127.5, 126.5, 123.5, 123.0, 122.5, 122.0, 119.0, 118.3, 83.0, 82.9, 77.7, 69.3, 69.2, 59.7, 56.4, 41.6, 36.8, 32.7, 32.6, 32.7, 28.3, 28.1, 28.0, 27.7, 25.8, 25.0, 24.8, 24.7, 13.5, 13.5. HRMS (ESI): calculated for $C_{21}H_{22}N_4OS$ $[M+H]^+$: 379.1587, found: 379.1591.

**NMR**

### 1.1. 1-(4-(6-Methylpyridin-2-yl)-6,7-dihydrothieno[3,2-*c*]pyridin-5(*4H*)-yl)ethan-1-one (3)

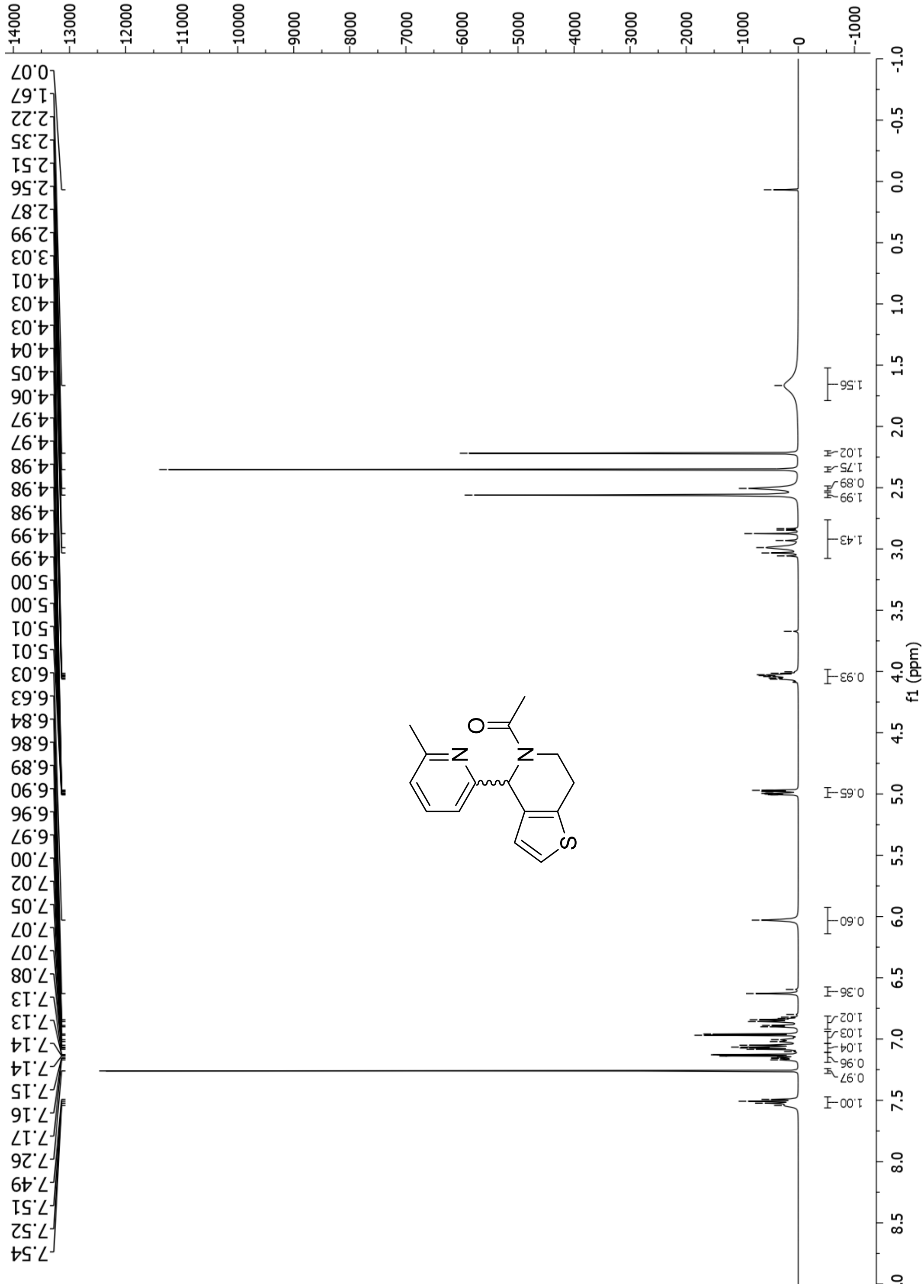

**Figure S8**. $^{1}$H NMR (500 MHz, $CDCl_3$, 298 K) of **3**.

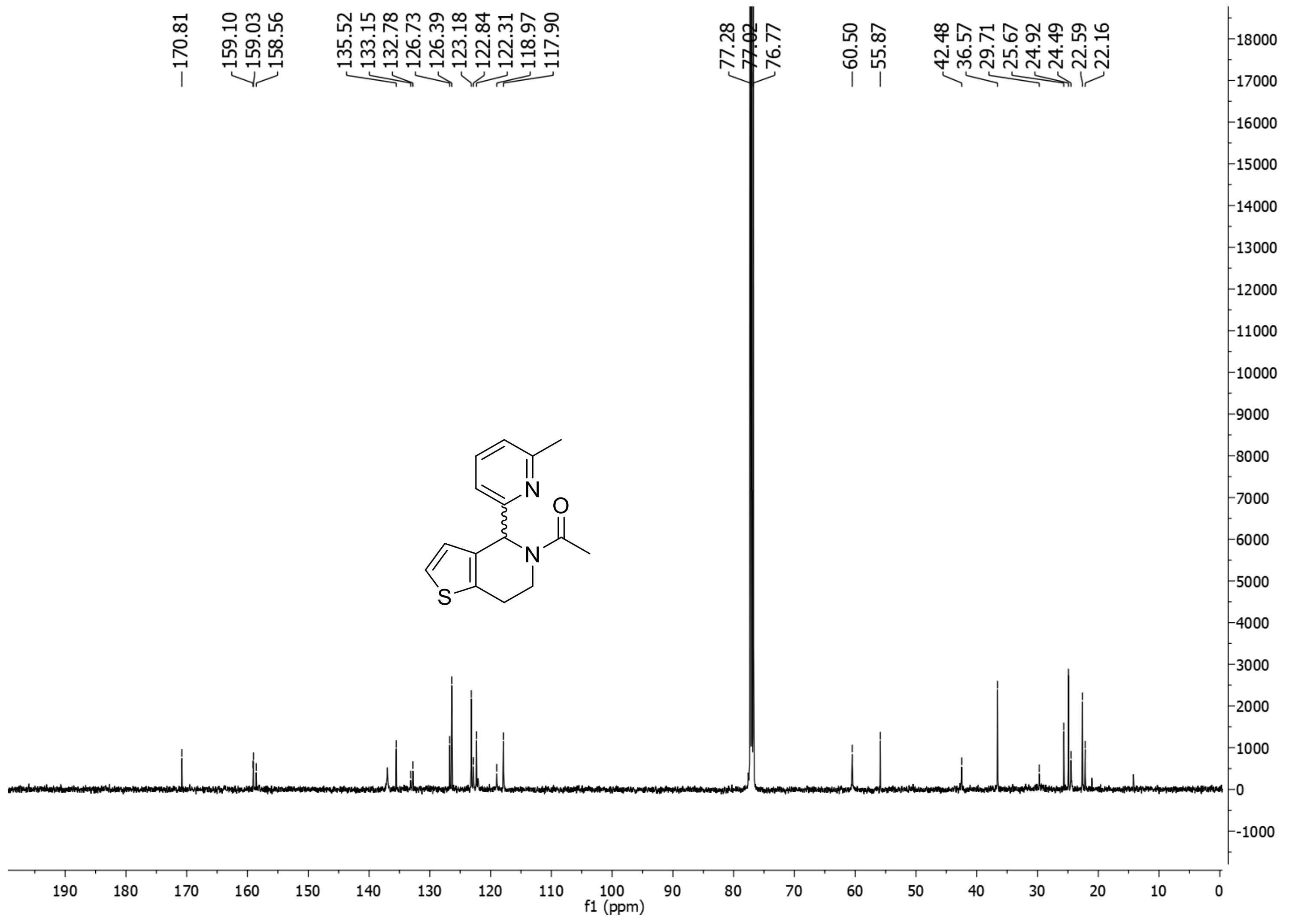

**Figure S9**. $^{13}$C NMR (500 MHz, $CDCl_3$, 298 K) of **3**.

### 1.2. 2-(S)-(2-Methylbutylamino)-1-[4-(6-methylpyridin-2-yl)-6,7-dihydro-*4H*-thieno[3,2-*c*]pyridin-5-yl]ethanone (4)

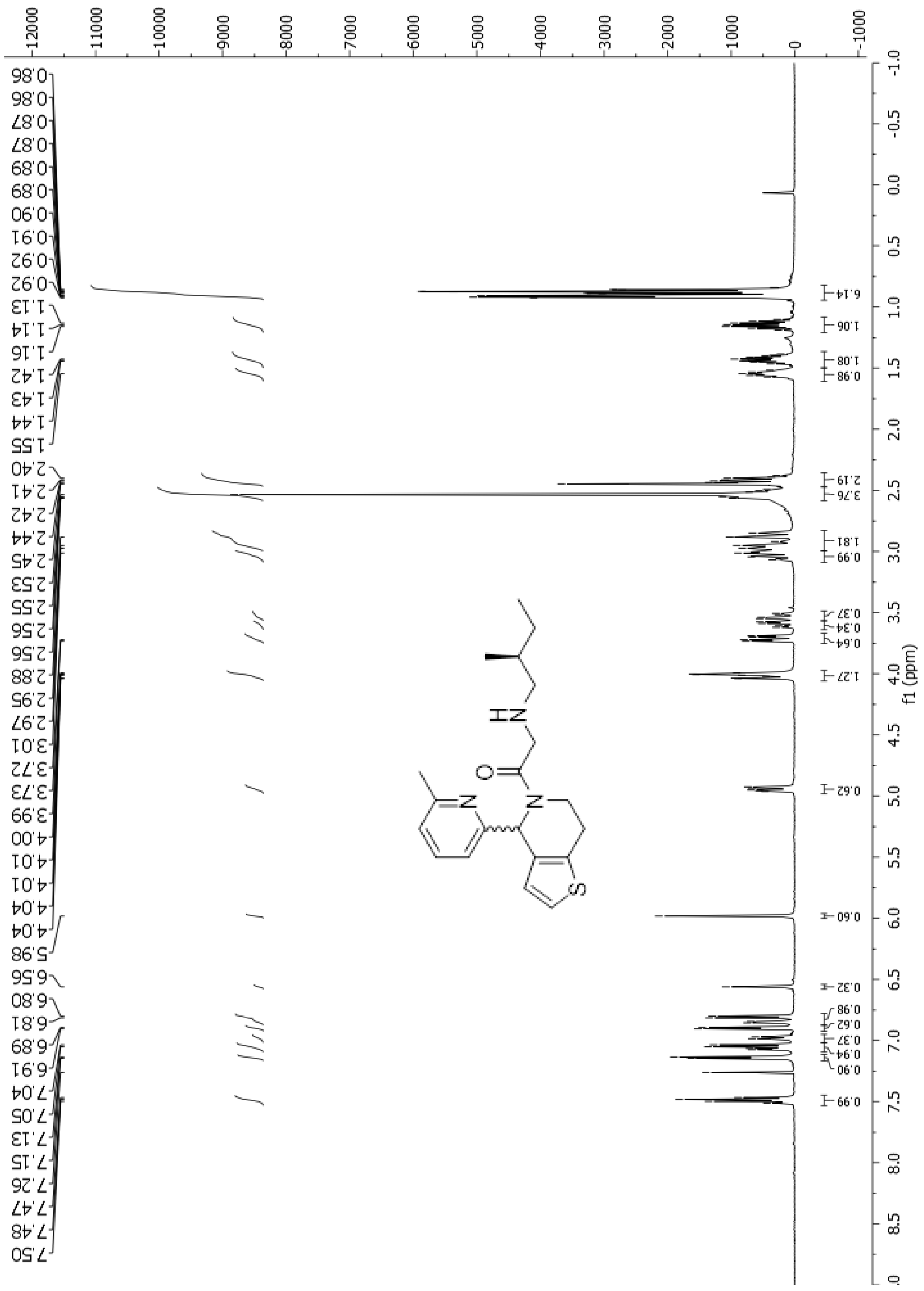

**Figure S10**. $^{1}H$ NMR (500 MHz, $CDCl_3$, 298 K) of **4**.

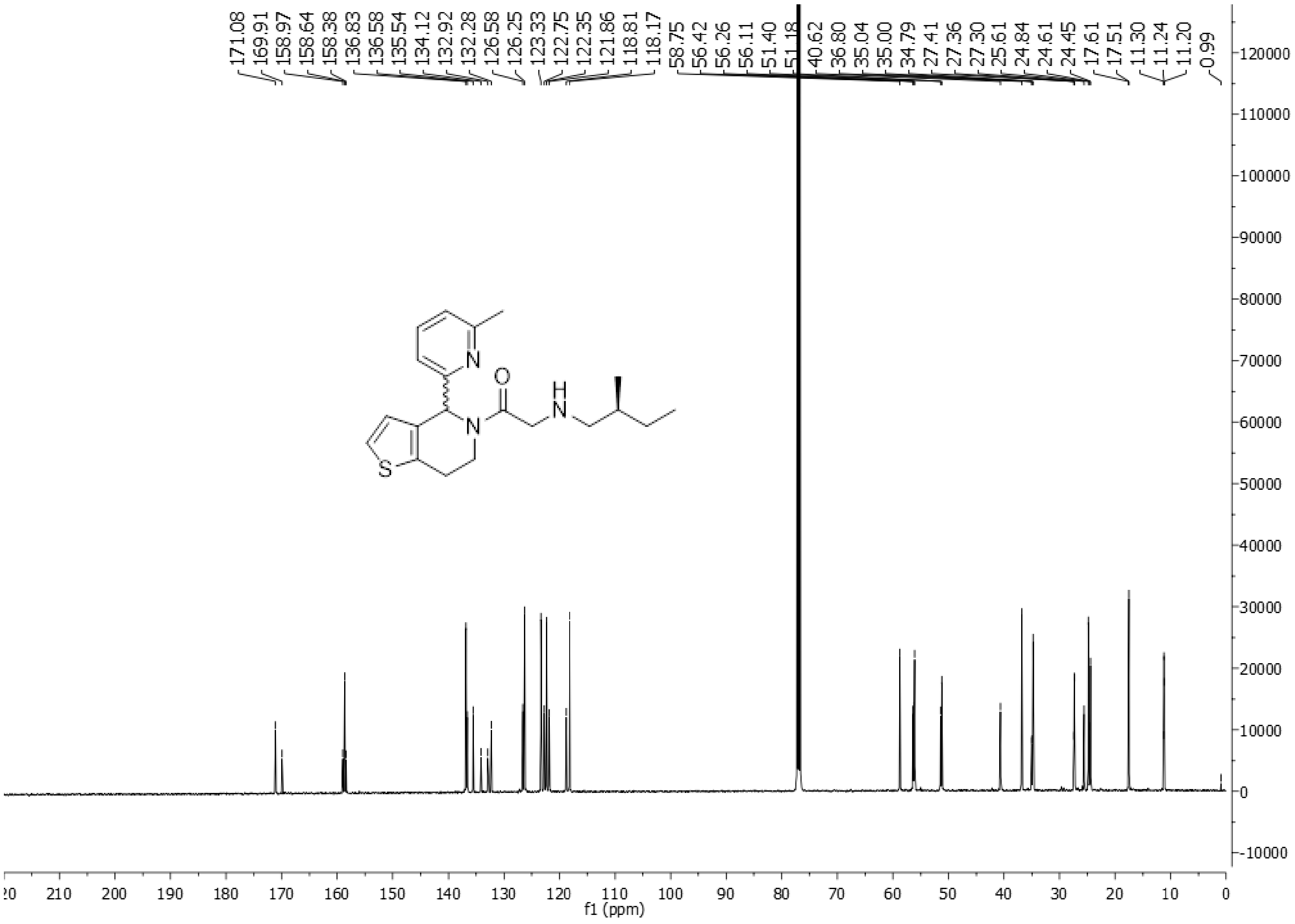

**Figure S11**. $^{13}$C NMR (500 MHz, $CDCl_3$, 298 K) of **4**.

### 1.3. 2-(Isopentylamino)-1-(4-(6-methylpyridin-2-yl)-6,7-dihydrothieno[3,2-*c*]pyridin-5(4*H*)-yl)ethan-1-one (5)

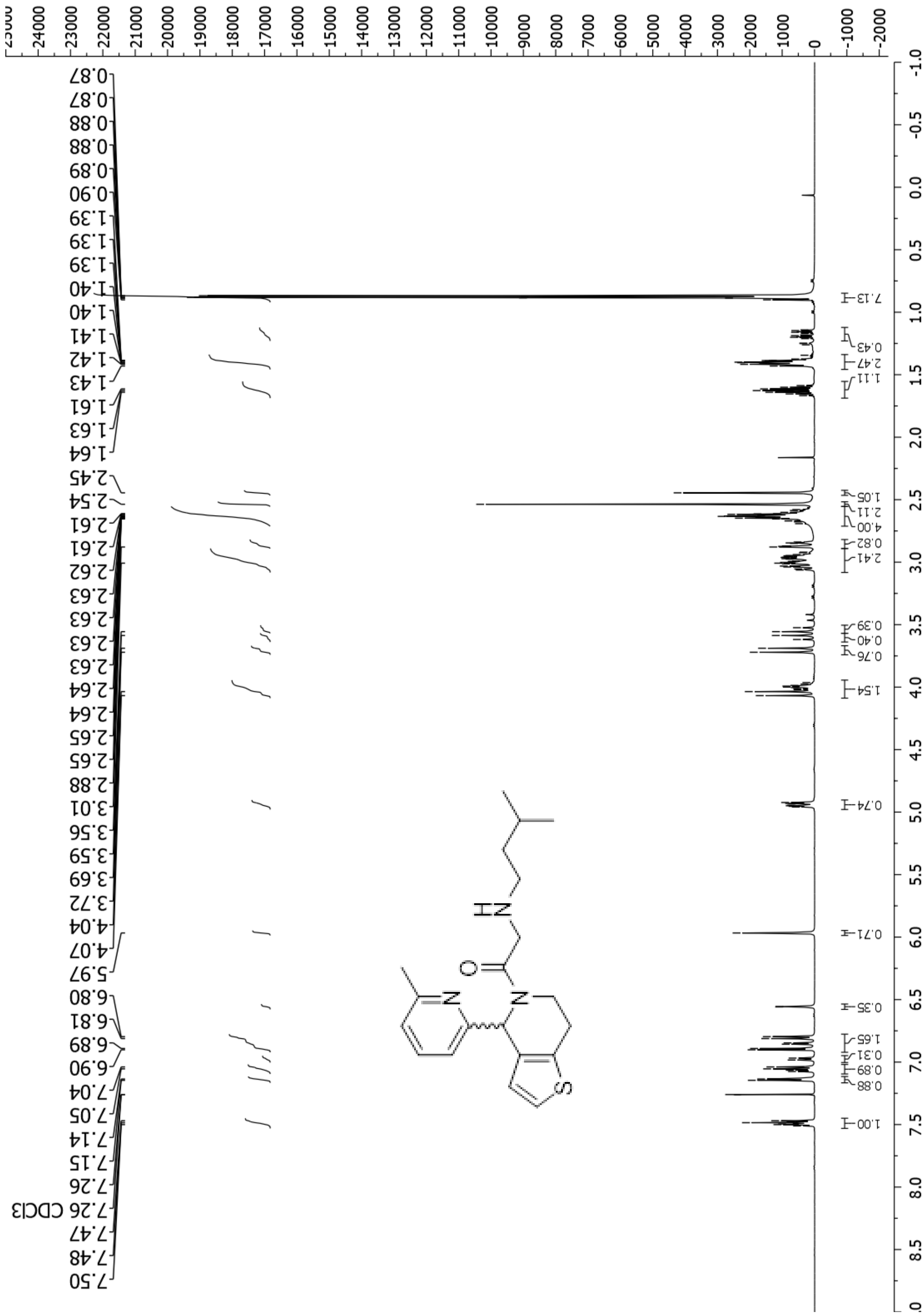

**Figure S12**. $^{1}$H NMR (500 MHz, $CDCl_3$, 298 K) of **5**.

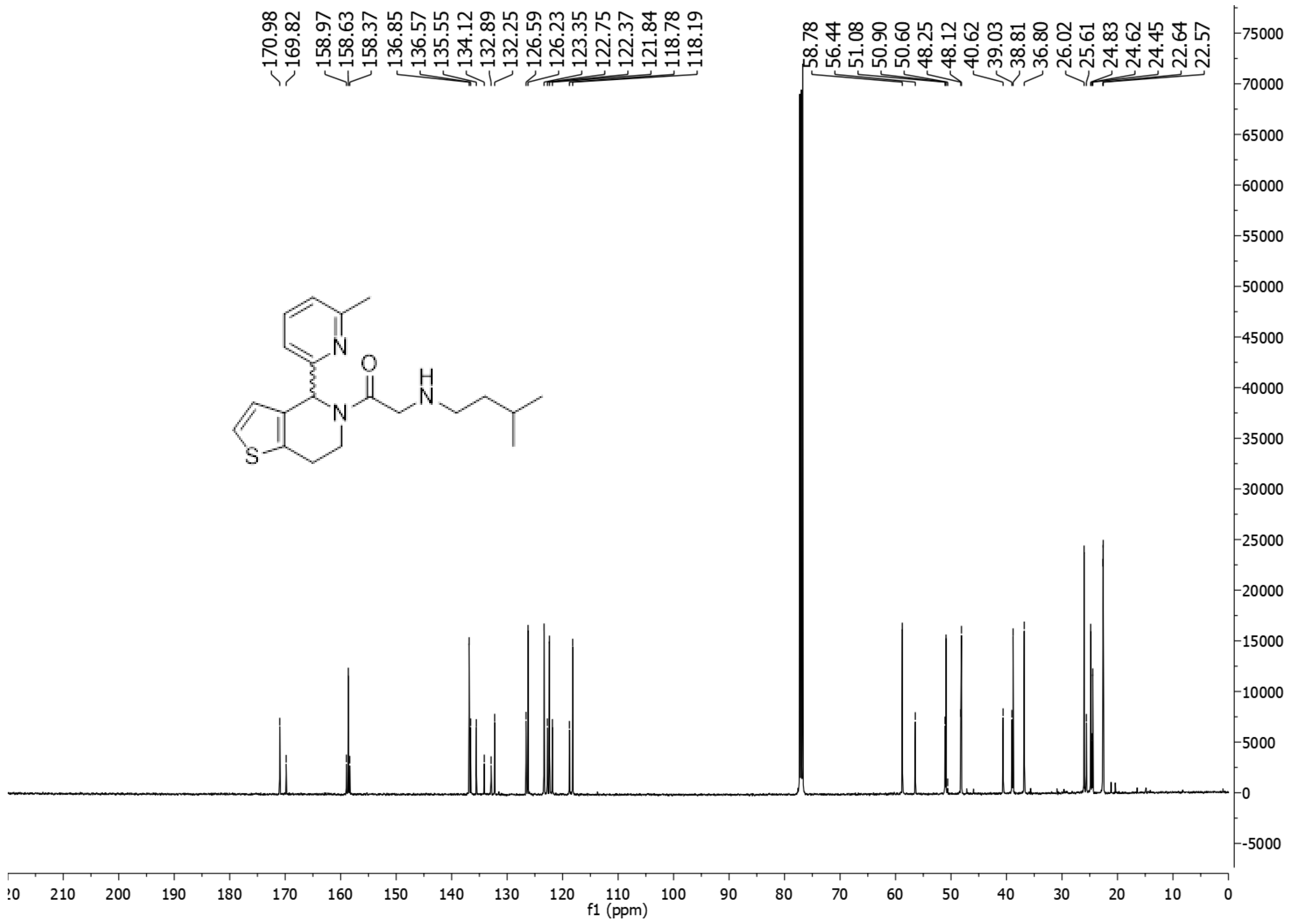

**Figure S13**. $^{13}$C NMR (500 MHz, $CDCl_3$, 298 K) of **5**.

### 1.4. 2-(2-Methylpropylamino)-1-[4-(6-methylpyridin-2-yl)-6,7-dihydro-4*H*-thieno[3,2-*c*]pyridin-5-yl]ethanone (6)

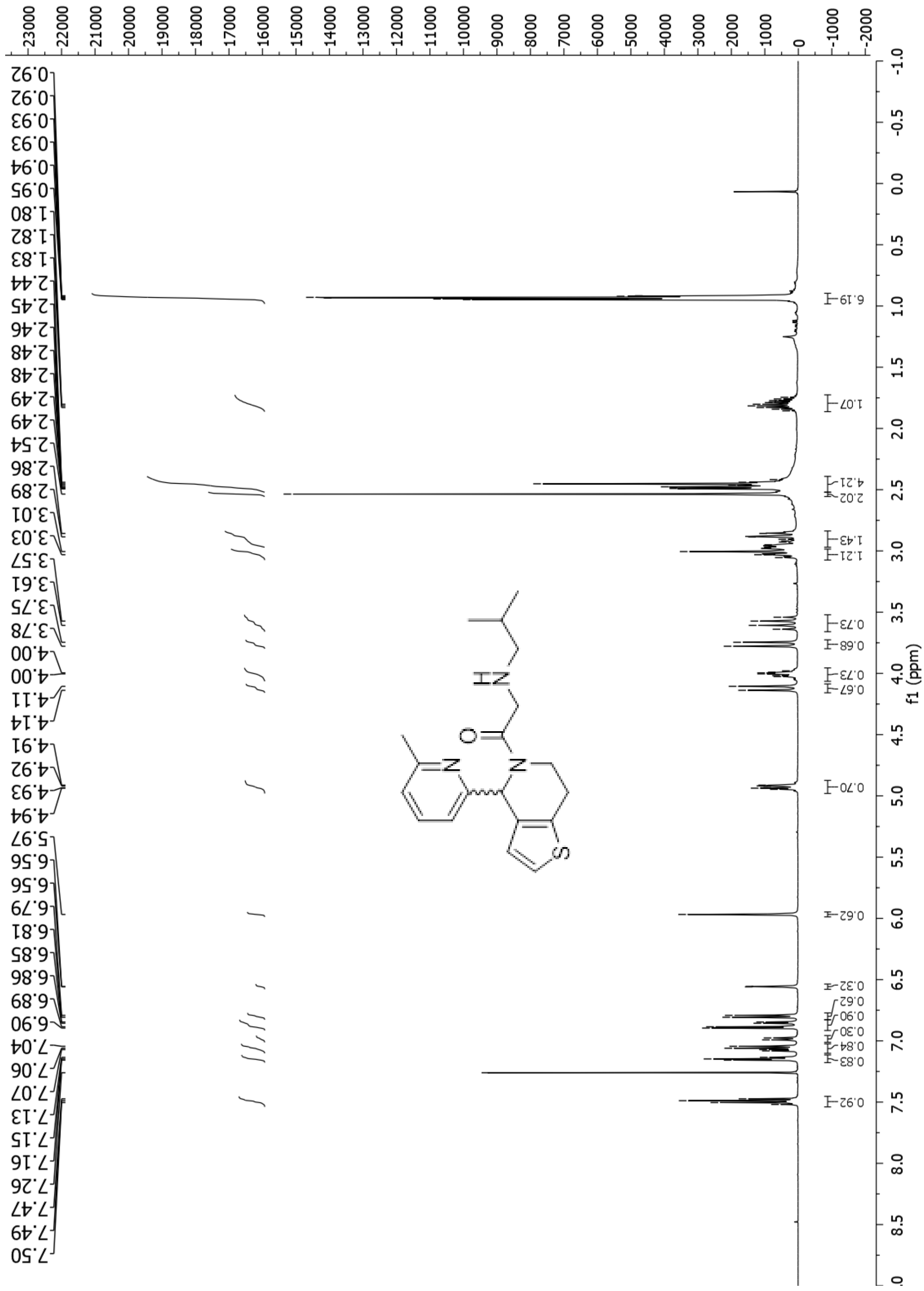

**Figure S14**. $^1$H NMR (500 MHz, $CDCl_3$, 298 K) of **6**.

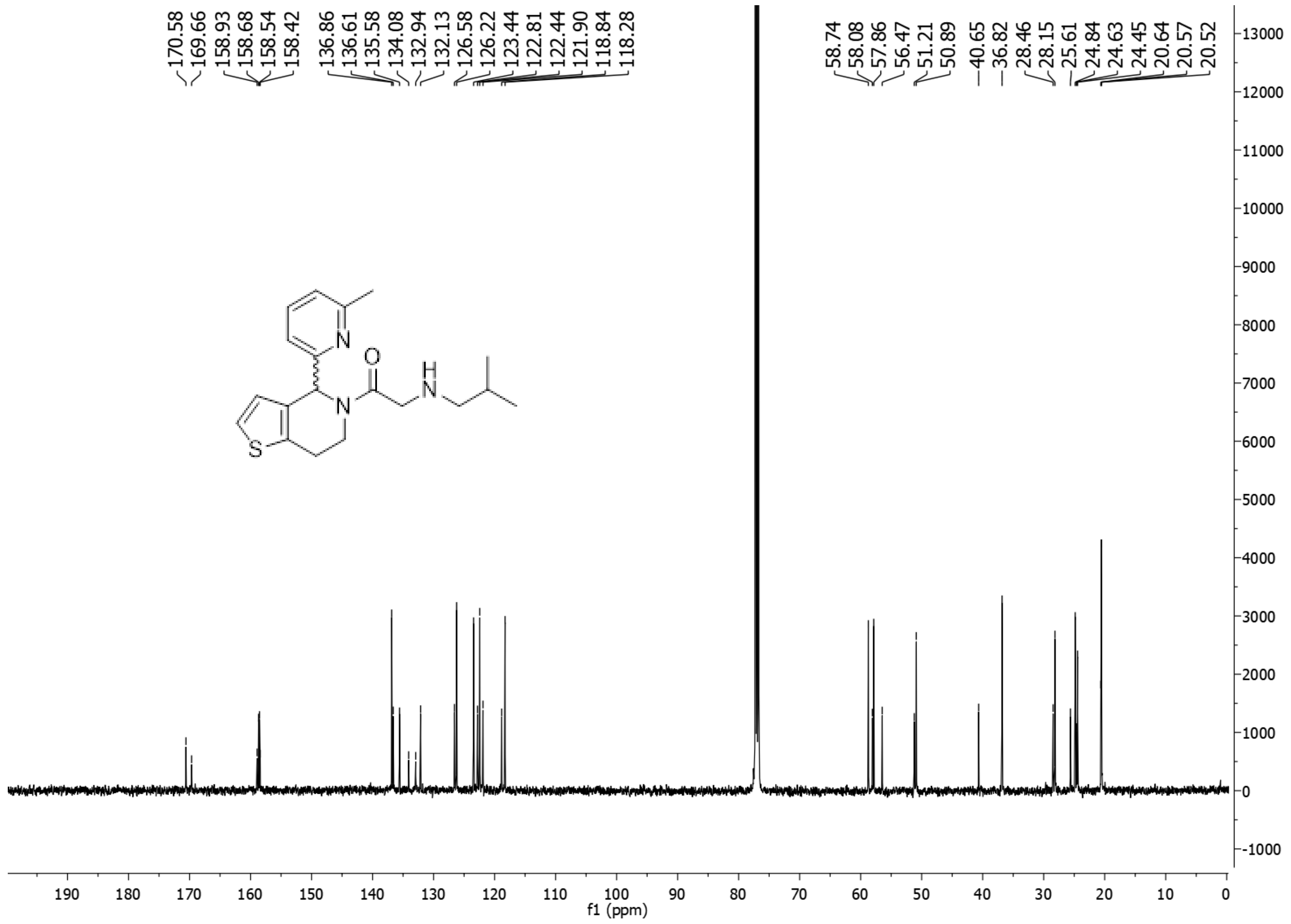

**Figure S15**. $^{13}C$ NMR (500 MHz, $CDCl_3$, 298 K) of **6**.

### 1.5. (-)-2-(2-Methylpropylamino)-1-[4-(6-methylpyridin-2-yl)-6,7-dihydro-4*H*-thieno[3,2-*c*]pyridin-5-yl]ethenone [(-)-6]

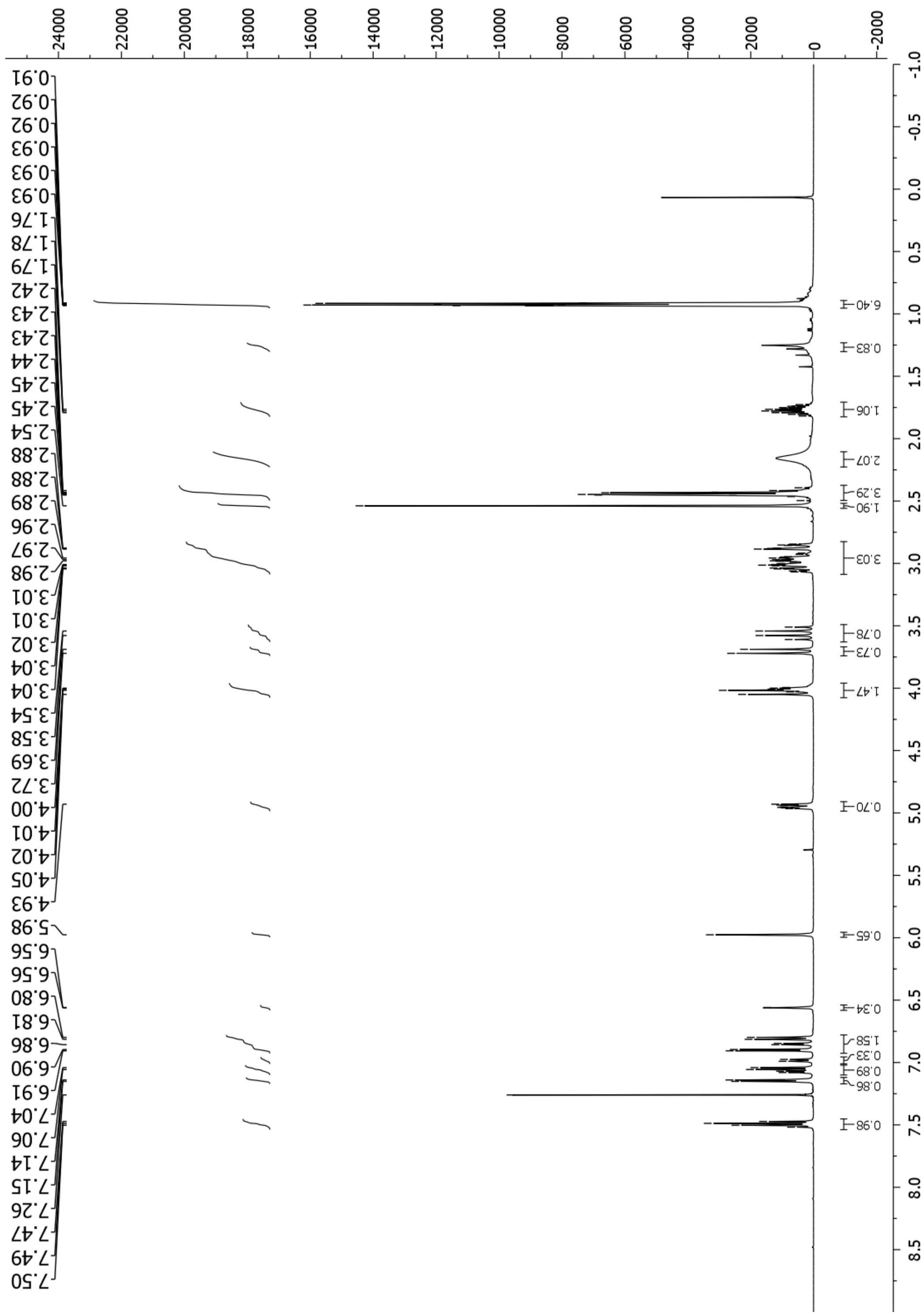

**Figure S16**. $^{1}$H NMR (500 MHz, $CDCl_3$, 298 K) of **(-)-6**.

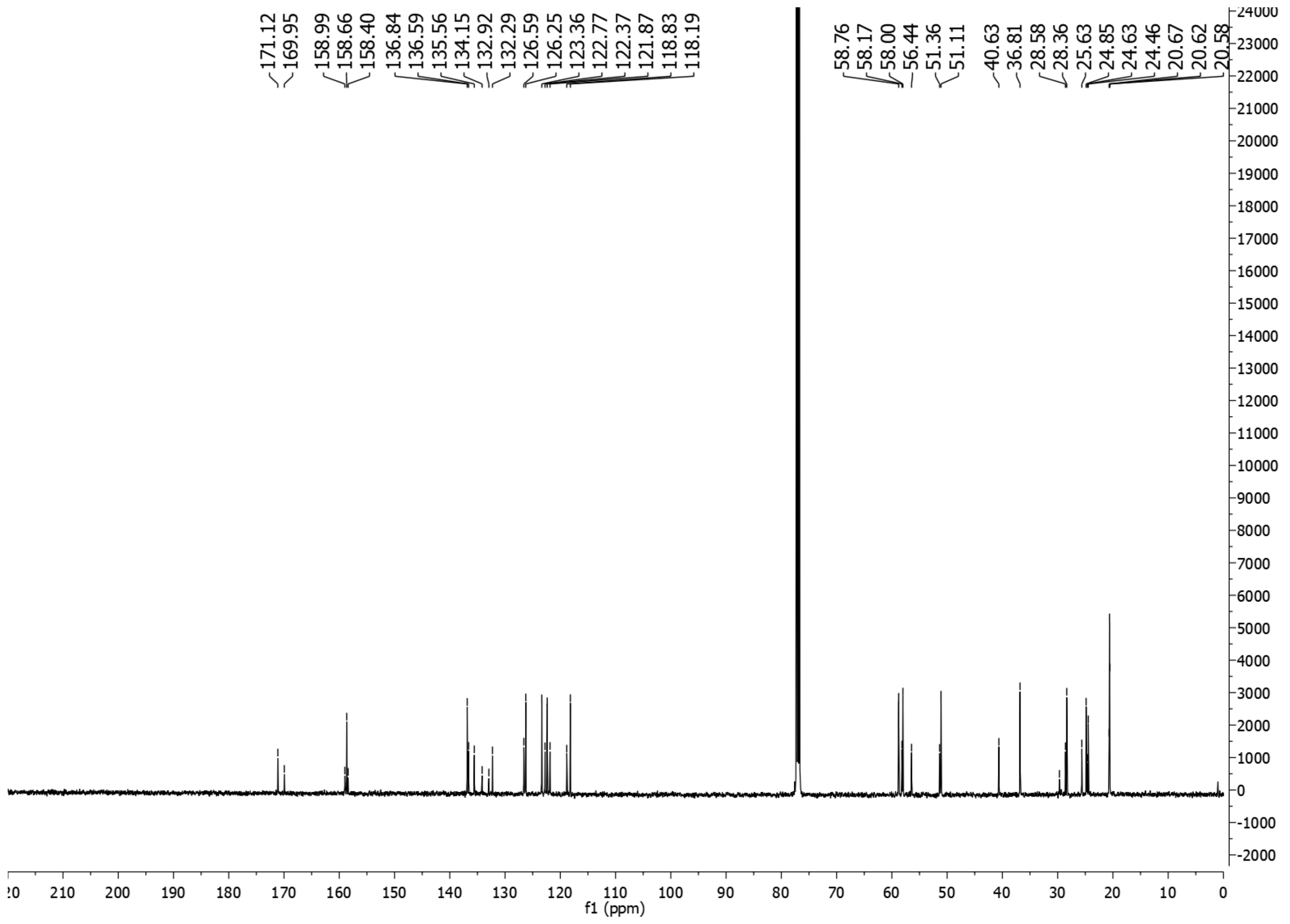

**Figure S17** $^{13}C$ NMR (500 MHz, $CDCl_3$, 298 K) of **(-)-6**.

### 1.6. (*+*)-2-(2-Methylpropylamino)-1-[4-(6-methylpyridin-2-yl)-6,7-dihydro-4*H*-thieno[3,2-*c*]pyridin-5-yl]ethanone [(+)-6, IMP-1575]

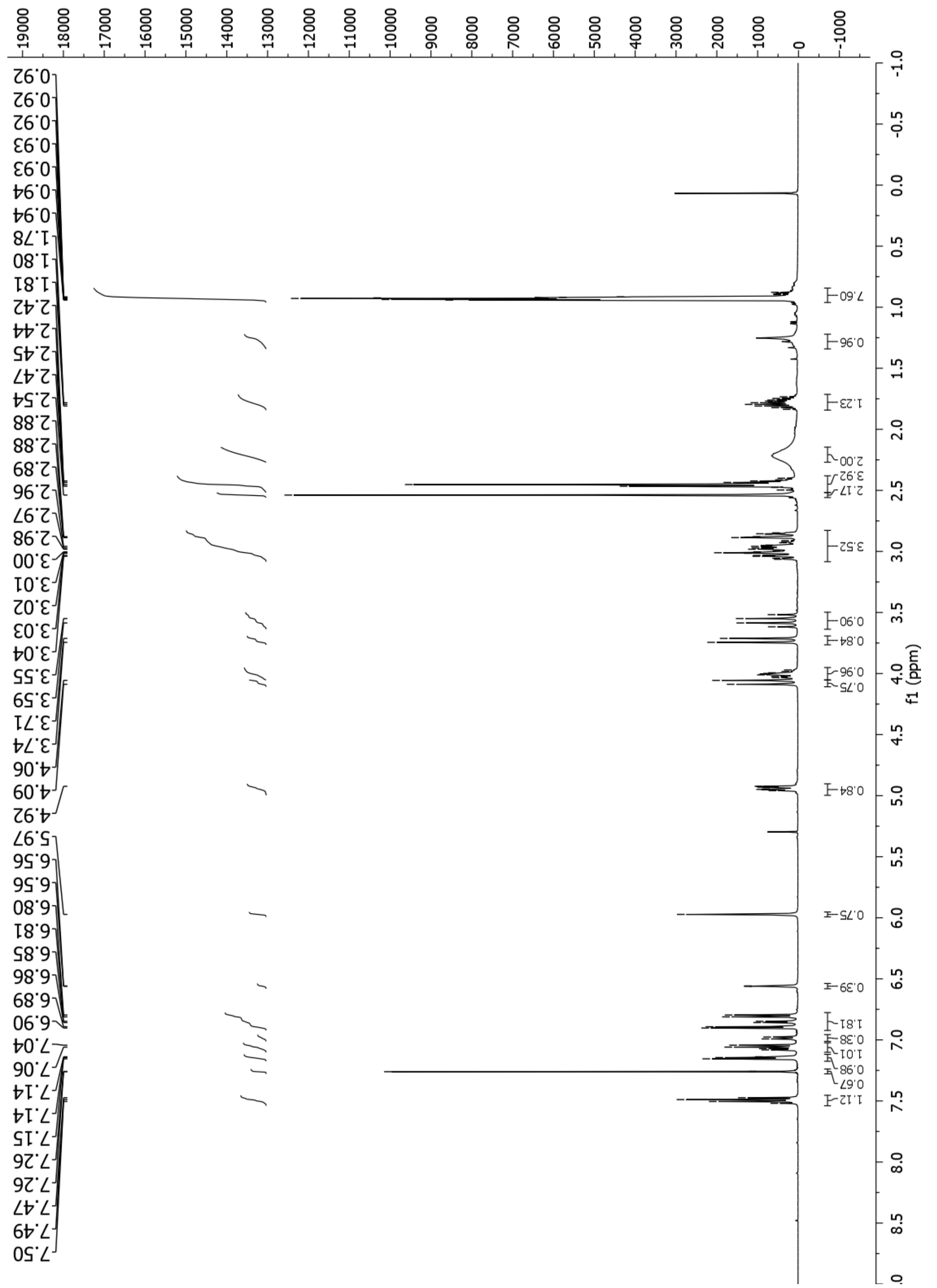

**Figure S18**. $^{1}H$ NMR (500 MHz, $CDCl_3$, 298 K) of (+)-**6** (**IMP-1575**).

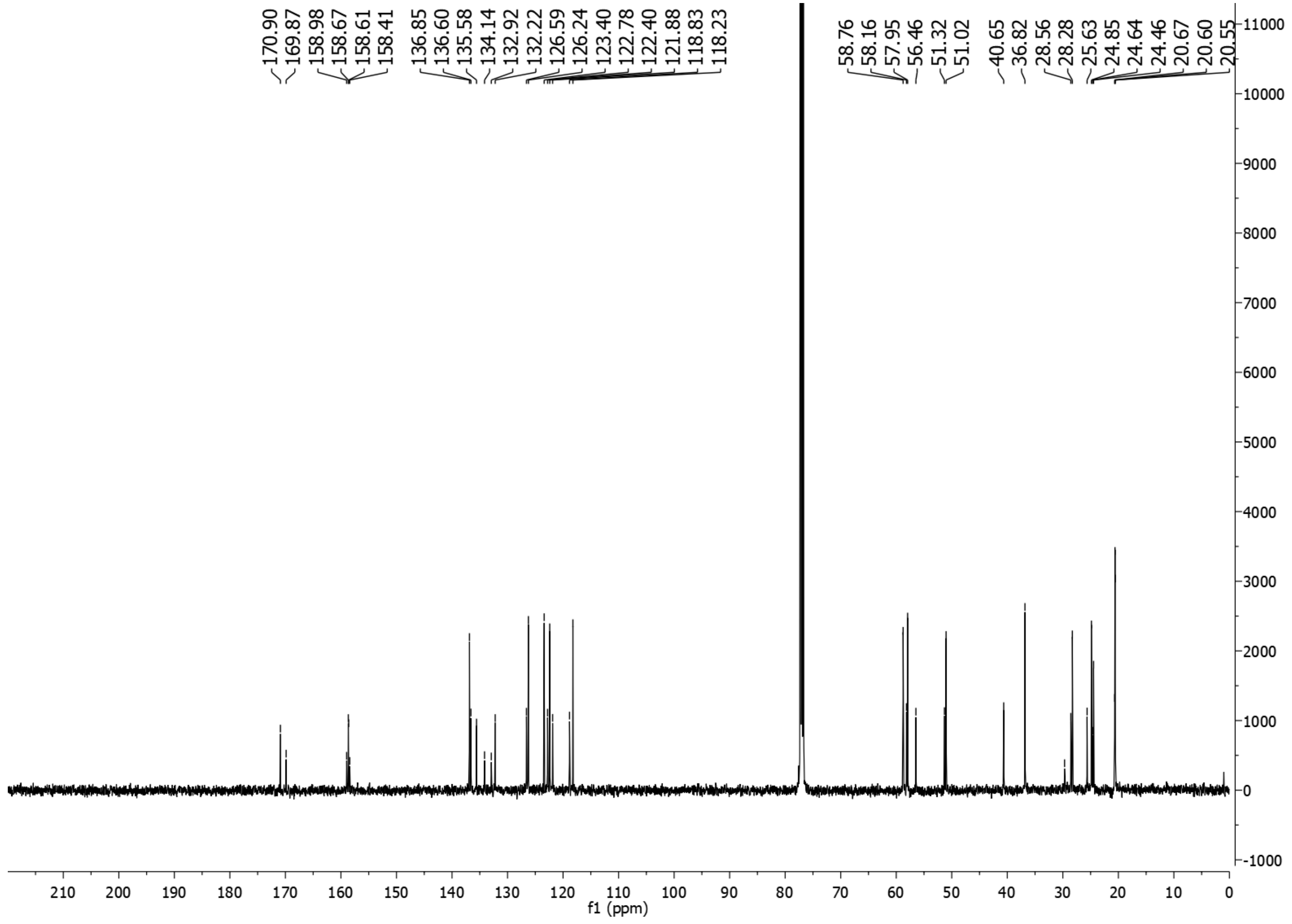

**Figure S19**. $^{13}$C NMR (500 MHz, $CDCl_3$, 298 K) of **(+)-6** (**IMP-1575**)

### 1.7. 2-(Isobutyl(methyl)amino)-1-(4-(6-methylpyridin-2-yl)-6,7-dihydrothieno[3,2-*c*]pyridin-5(4*H*)-yl)ethan-1-one (7)

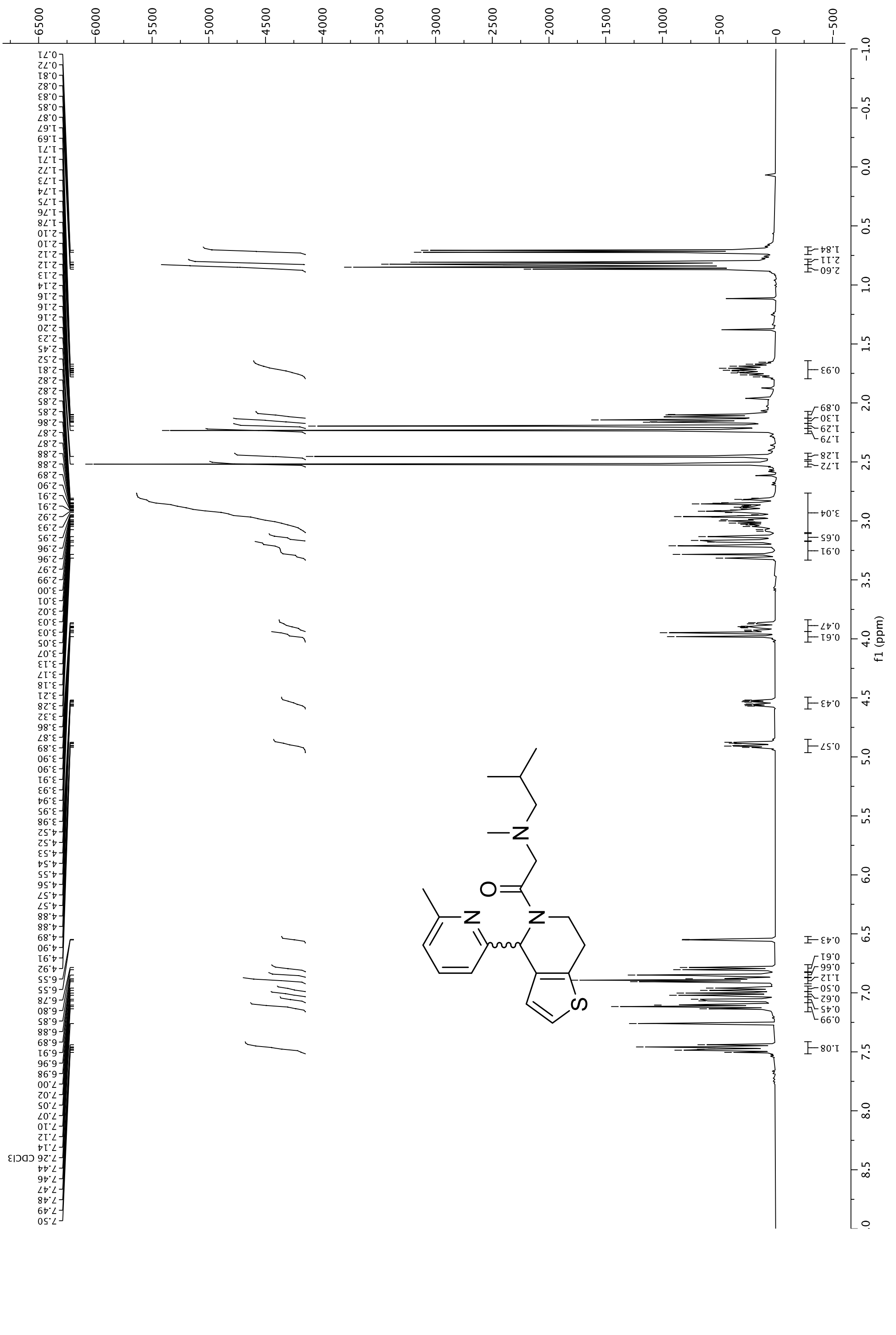

**Figure S20**. $^{1}H$ NMR (500 MHz, $CDCl_3$, 298 K) of **7**.

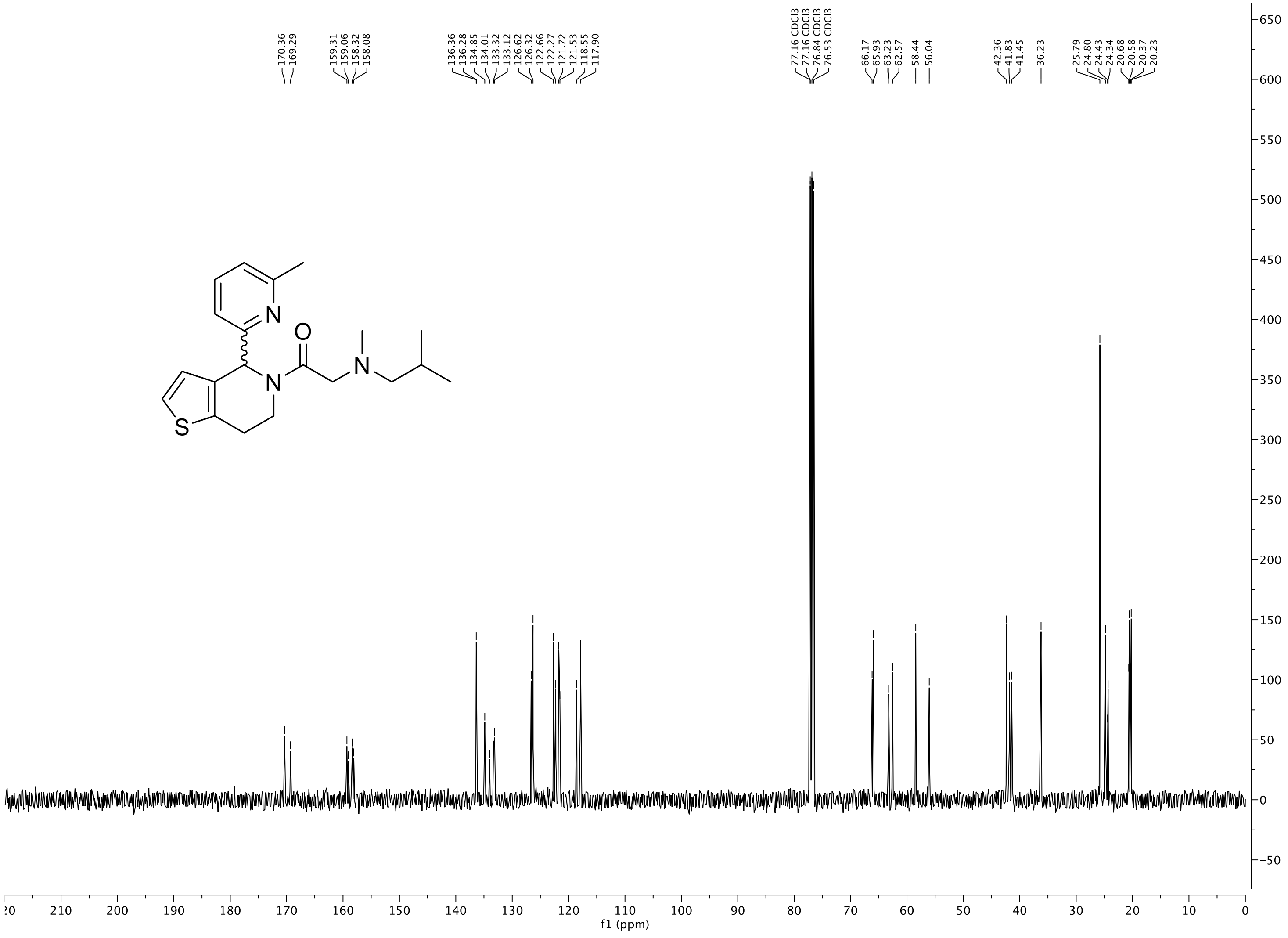

**Figure S21**. $^{13}C$ NMR (500 MHz, $CDCl_3$, 298 K) of **7**.

### 1.8. 2-(3-Methylpiperidin-1-yl)-1-(4-(6-methylpyridin-2-yl)-6,7-dihydrothieno[3,2-*c*]pyridin-5(4*H*)-yl)ethan-1-one (8)

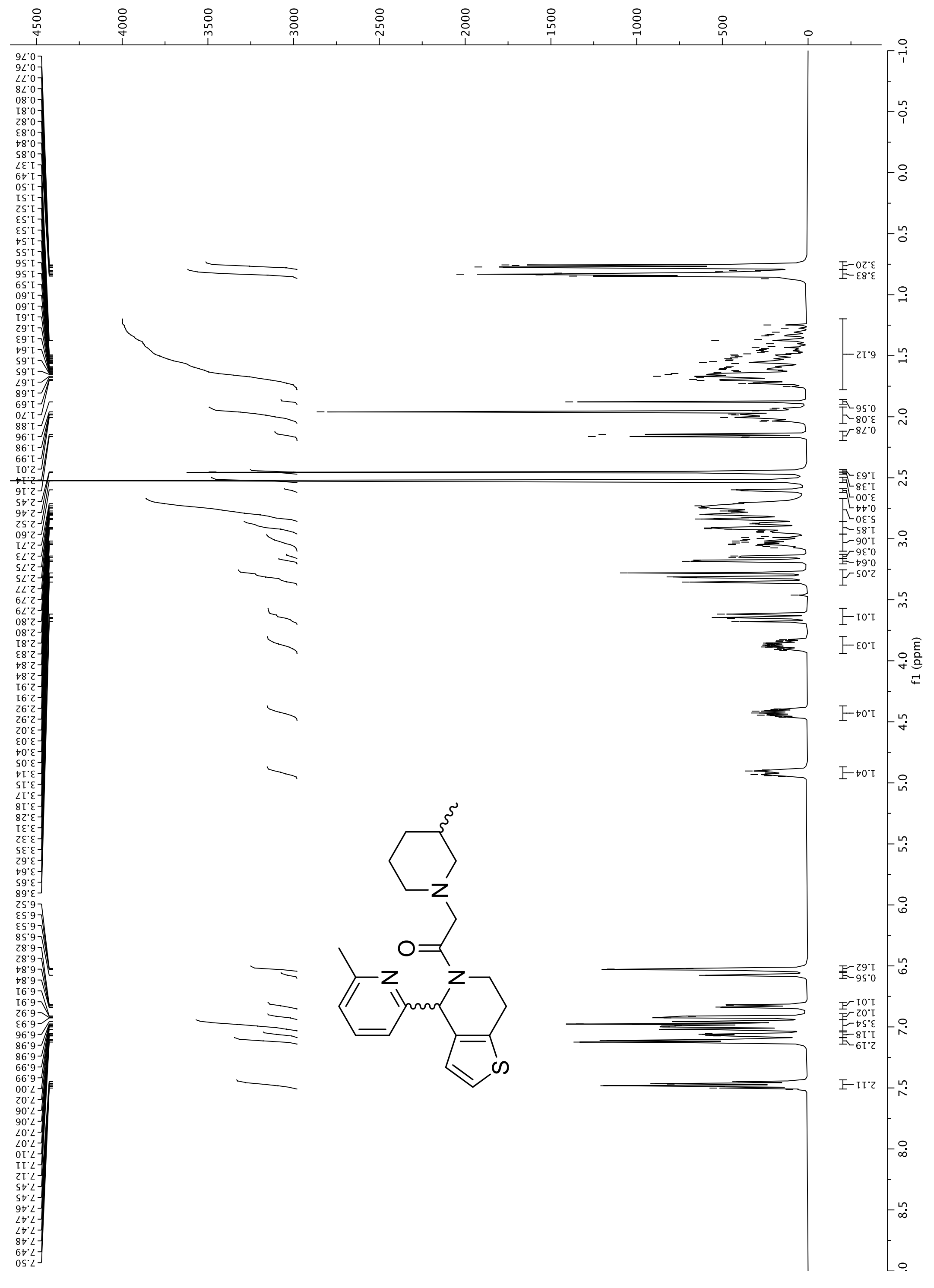

**Figure S22**. $^{1}H$ NMR (500 MHz, $CDCl_3$, 298 K) of **8**.

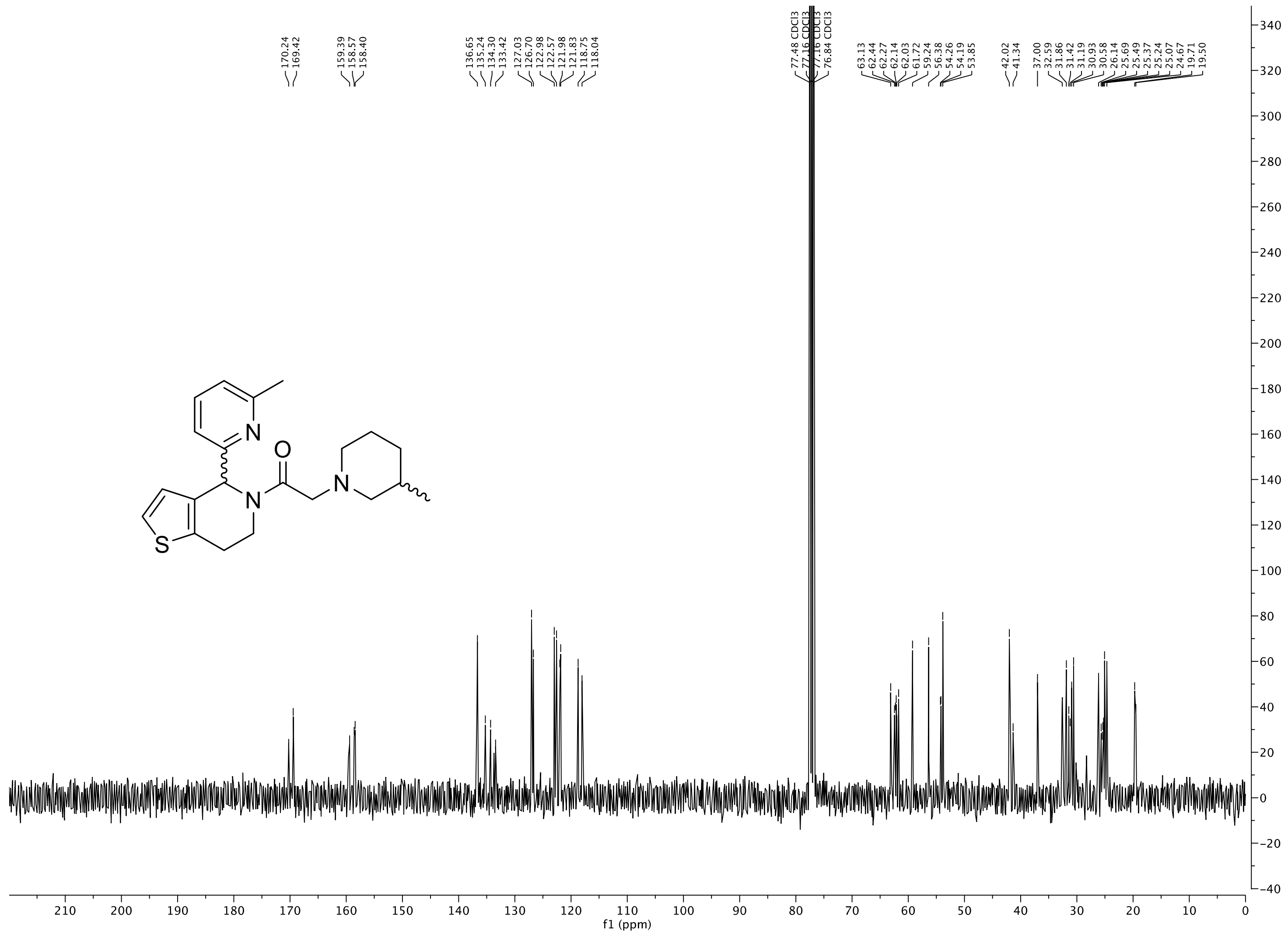

**Figure S23**. $^{13}$C NMR (500 MHz, $CDCl_3$, 298 K) of **8**.

### 1.9. *N*-isopentyl-4-(6-methylpyridin-2-yl)-6,7-dihydrothieno[3,2-*c*]pyridine-5(4*H*)-carboxamide (9)

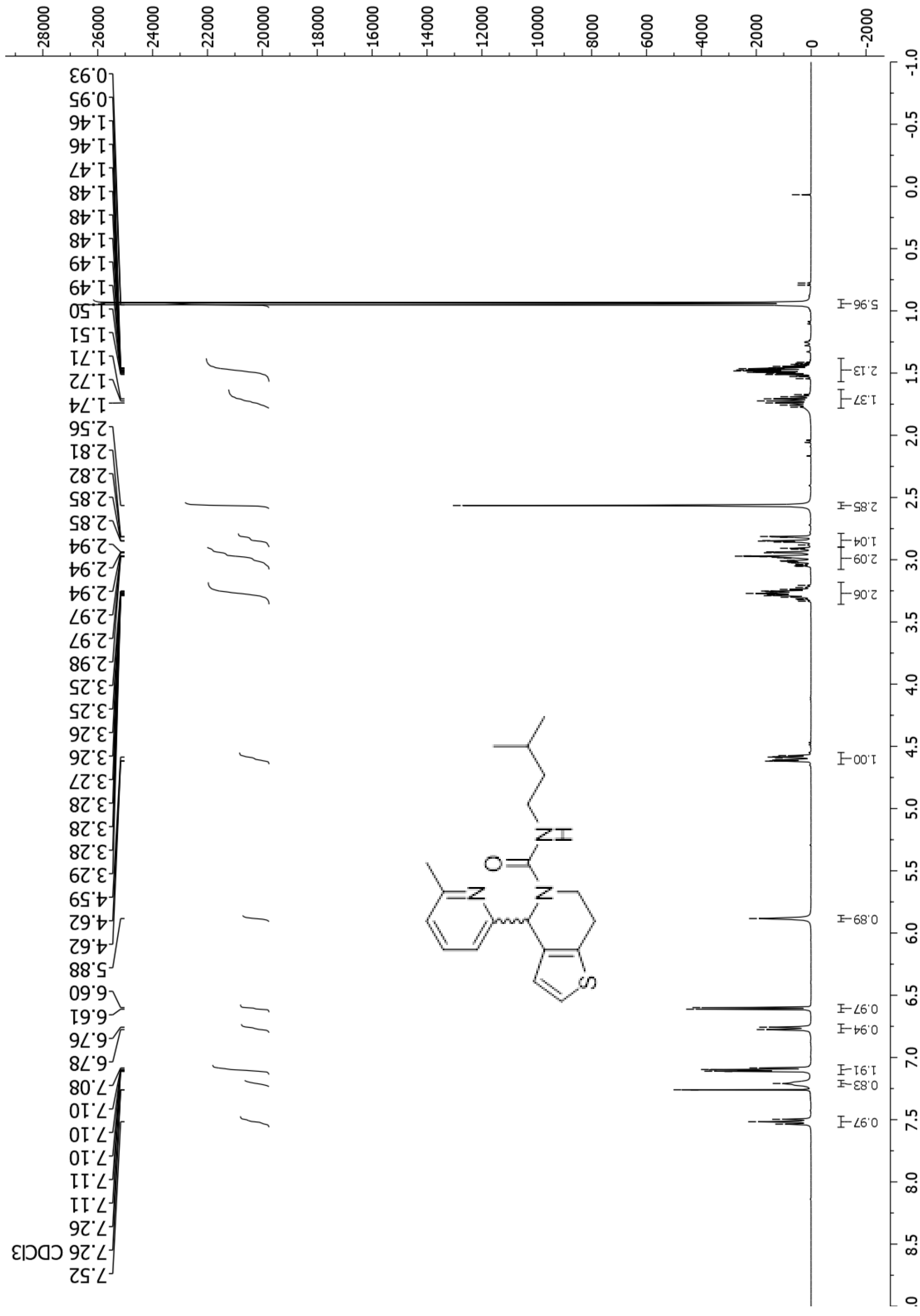

**Figure S24**. $^{1}$H NMR (400 MHz, $CDCl_3$, 298 K) of **9**.

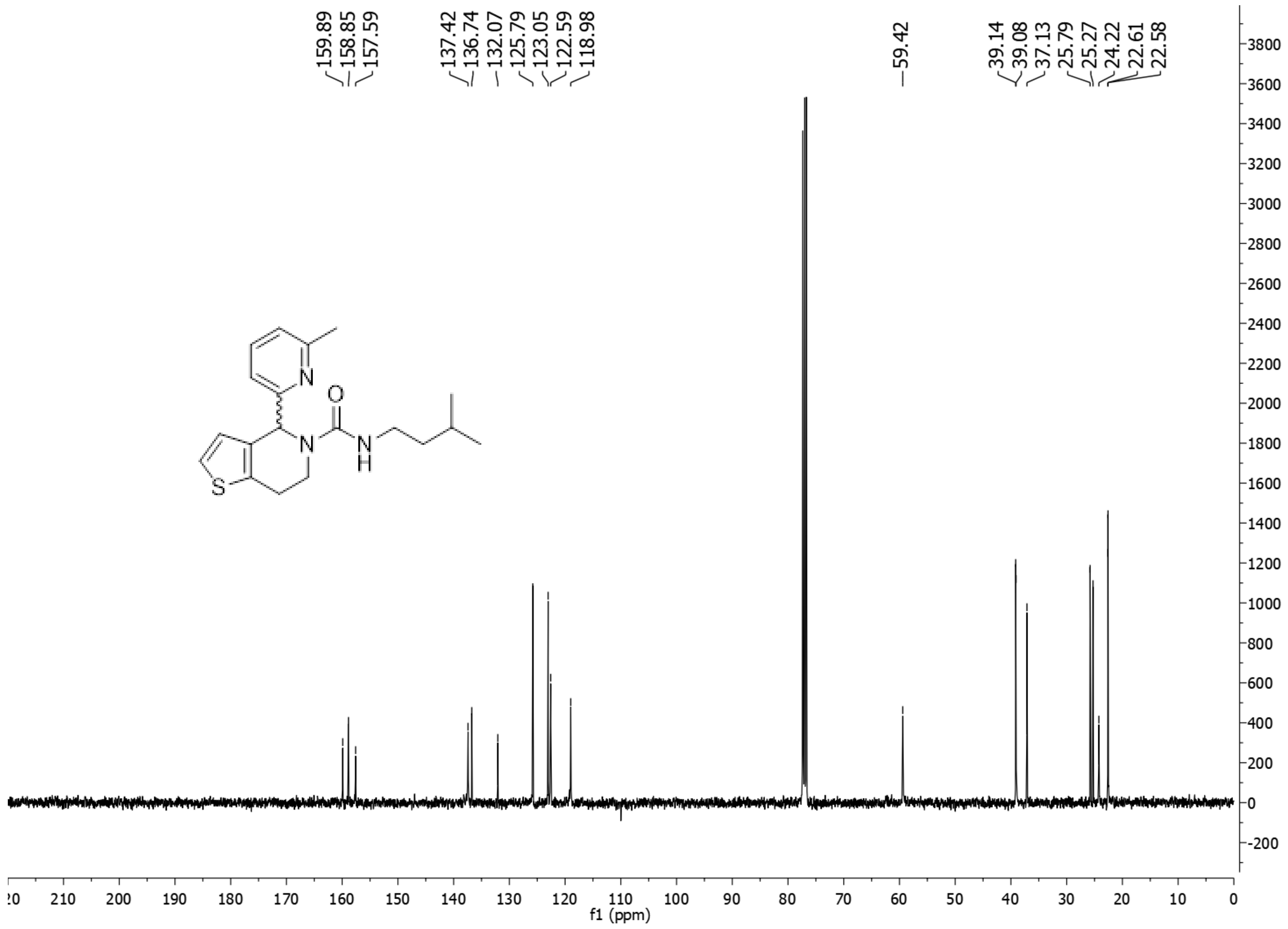

**Figure S25**. $^{13}C$ NMR (400 MHz, $CDCl_3$, 298 K) of **9**.

### 1.10. 3-(3-(But-3-yn-1-yl)-*3H*-diazirin-3-yl)-1-(4-(6-methylpyridin-2-yl)-6,7-dihydrothieno[3,2-*c*]pyridin-5(*4H*)-yl)propan-1-one (10)

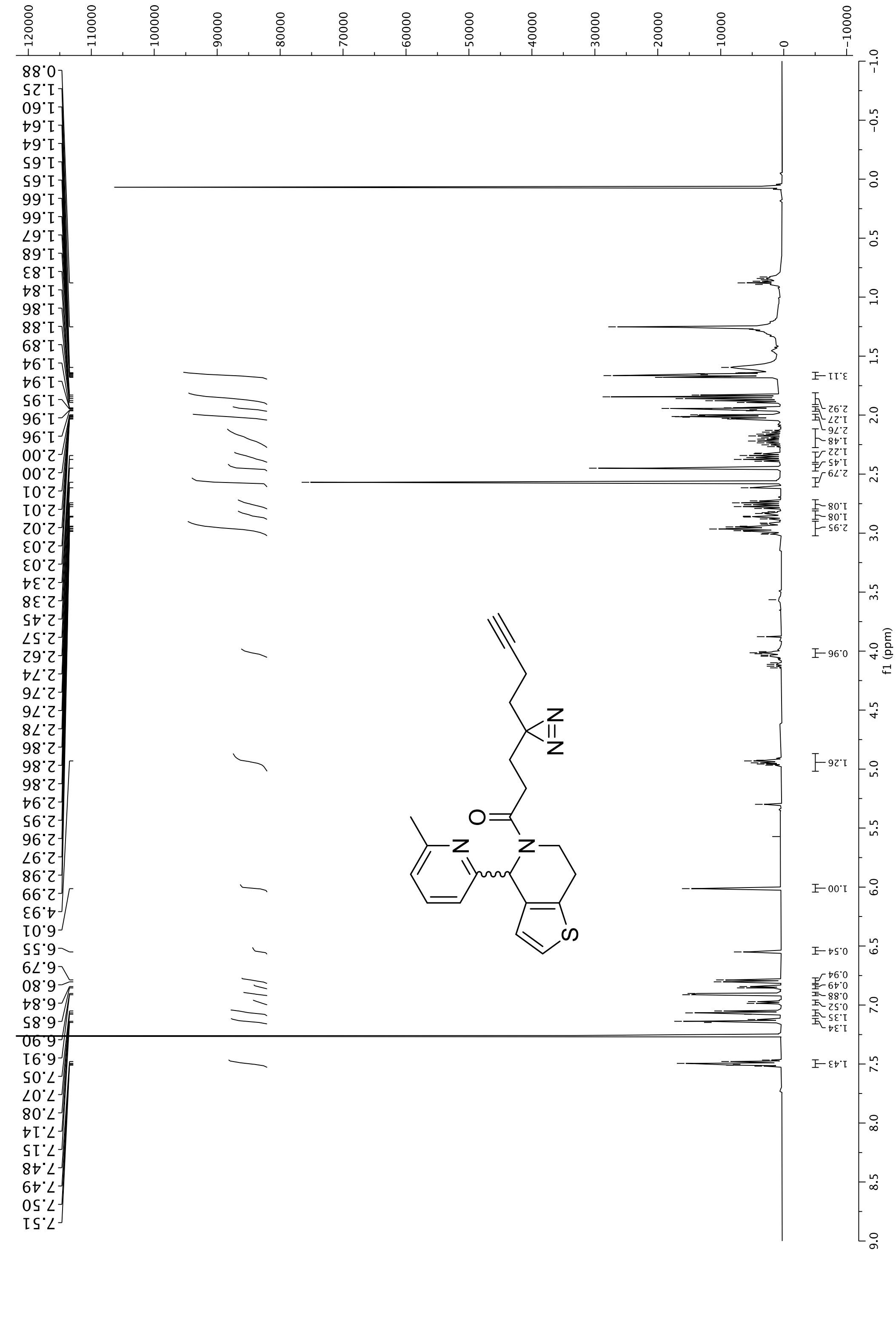

**Figure S26**. $^1$H NMR (500 MHz, $CDCl_3$, 298 K) of **10**.

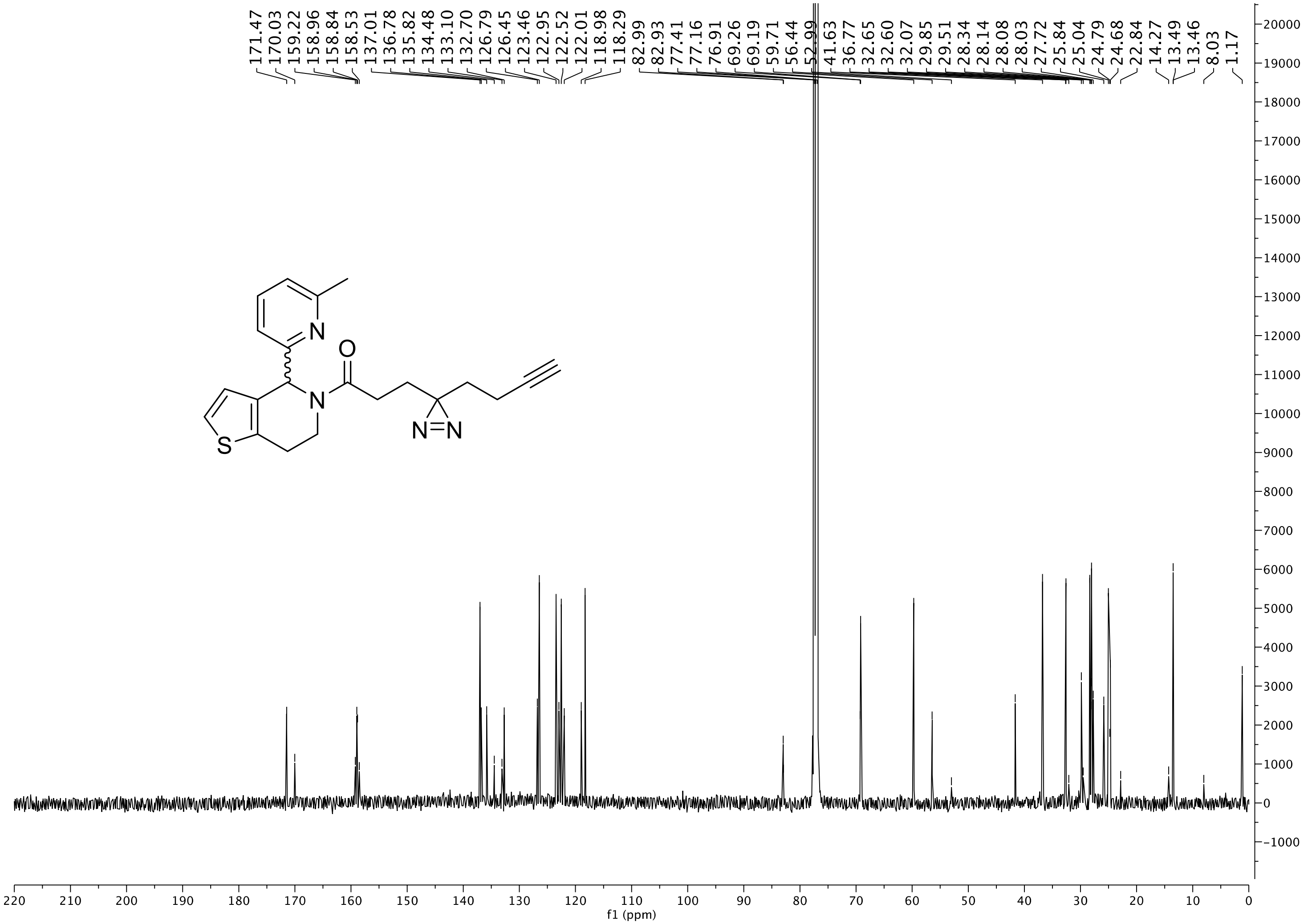

**Figure S27**. $^{13}$C NMR (500 MHz, $CDCl_3$, 298 K) of **11**.
